# Antifungal biosynthesis by root-associated *Streptomyces* and *Pseudomonas* is elicited upon plant colonization

**DOI:** 10.1101/2025.04.26.650784

**Authors:** Wendalina Tigani, Jack G. Ganley, Chao Du, Somayah S. Elsayed, Emtinan Diab, Paolo Innocenti, Iulian Rimboi, Victor Carrion-Bravo, Nathaniel I. Martin, Mohammad R. Seyedsayamdost, Jos M. Raaijmakers, Gilles P. van Wezel

## Abstract

Plants are colonized by a diverse microbiome, with microorganisms residing inside and outside of plant tissues. Plants can harness the protective traits of their microbial inhabitants to ward off insect pests and fungal pathogens. However, current understanding of the role of commensal interactions on activating the desired microbial genomic traits remains limited. Here we show that biosynthesis of the antifungal 2,5-dihydro-L-phenylalanine (DHP) by the endophytic *Streptomyces* sp. PG2 is strongly induced upon colonization of *Arabidopsis thaliana*. DHP production protects the plant from infection by the fungal root pathogen *Rhizoctonia solani*, both *in vitro* and *in vivo*.. We identified the DHP biosynthetic gene cluster (BGC) and showed that heterologous expression of the BGC in the DHP non-producer *Streptomyces coelicolor* also conferred plant-inducible DHP production. The BGC was also found in plant-associated Gram-negative bacteria, and in *Pseudomonas syringae* FF5 we again observed strongly enhanced DHP production upon plant colonization. An ecology-inspired elicitor screen showed that L-valine and brassinosteroid hormones elicit DHP biosynthesis in the plant-beneficial *Streptomyces* sp. PG2, while L-valine also elicited DHP biosynthesis in *S. coelicolor*. *In vivo* experiments confirmed the stimulation of antifungal activity in *Streptomyces* sp. PG2 by L-valine, while brassinolide mutant plants showed reduced DHP induction. Conversely, neither L-valine nor brassinolide elicited the expression of the DHP BGC in the pathogenic *P. syringae*, revealing important divergence in the responses to plant signaling, which may reflect selectivity in how endosymbionts and pathogens respond to host cues. Collectively, our data demonstrate that plant colonization can elicit the biosynthetic potential of root-associated microbes, thereby enhancing plant resilience.

## INTRODUCTION

Plants are home to taxonomically and functionally diverse communities of bacteria, fungi, protists, nematodes and viruses, collectively referred to as the plant microbiome. These plant-associated microorganisms can provide various beneficial functions for their host by enhancing plant growth, facilitating nutrient acquisition, and bolstering resistance against pathogens^1^. Plants secrete compounds that actively shape the microbiome of the rhizosphere^2^. In return, plant growth-promoting bacteria (PGPB) suppress disease caused by microbial pathogens^3, 4^ through several mechanisms, including competition for nutrients and iron, the production of lytic enzymes and the biosynthesis of bioactive molecules such as antibiotics or antifungals^5, 6^. Members of the genus *Streptomyces*, the largest genus within the Actinobacteria, produce a wide range of secondary metabolites with antibacterial, antifungal, antiviral, anticancer or nematicidal activities^7, 8^. Streptomycetes are prominent members of the plant microbiome and play an important role in enhancing plant tolerance to biotic stresses^9–11^. However, most knowledge of their bioactivities comes from studies conducted under controlled laboratory conditions, whereas genomic analyses point to a much greater biosynthetic potential^11–13^. A better understanding of *Streptomyces* ecology, their interactions with the plant host and the underlying chemistry is needed to uncover and harness their largely untapped functional potential^14^.

Every step of the association between plants and microorganisms is tightly regulated by the exchange of specific signaling molecules. Plant-associated microorganisms perceive signals released by the plant in the root or shoot exudates and actively move toward the plant^15, 16^. Root exudate constituents are key for recruiting specific microorganisms from the diverse bulk soil microbial reservoir, resulting in distinct signatures of microbiome assembly for a given plant species or even cultivars within a plant species^17^. Plant signals can also trigger bacterial attachment and biofilm formation on the root surface^18^. Following colonization, plants may undergo metabolic changes, with the rhizosphere microbiome impacting root exudation profile^19^, plant secondary metabolism^20^ and immunity (Induced Systemic Resistance, ISR)^6^. However, the reciprocal effects of the host plant on the chemistry and activities of the recruited microorganisms remain largely elusive.

The “cry for help” hypothesis^21^ entails that plants harness their microbiome to protect themselves against pathogens by eliciting the biosynthesis of bacterial secondary metabolites. Indeed, plant hormones have been shown to elicit the biosynthesis of bioactive compounds^11, 22^. We recently demonstrated *in vitro* that jasmonic acid (JA) and methyl jasmonate (MeJA) alter growth and trigger antibiotic production in streptomycetes^23^, while the ubiquitous plant metabolite catechol also elicits natural product biosynthesis in these bacteria^24^. However, how plants ensure that beneficial microbes in their microbiome produce disease-suppressive natural products in the rhizosphere and especially at the time of infection, remains a major question to be resolved.

We sought to investigate the ability of root-associated bacteria to interact with plants *in vivo* and protect against fungal infections. To do so, we studied the ability of root-associated *Streptomyces* to protect *Arabidopsis thaliana* against infection by the fungal root pathogen *Rhizoctonia solani*. Using genetics, phylogenomics, metabolomics, and high-throughput elicitor screening (HiTES), we discovered that *A. thaliana* metabolites, specifically L-valine and brassinosteroid hormones, strongly elicit the biosynthesis of the antifungal 2,5-dihydro-L-phenylalanine (DHP) in the endophytic *Streptomyce*s sp. PG2. DHP inhibited hyphal growth of *R. solani* and suppressed its infection of plants, both *in vitro* and *in vivo*. Interestingly, the DHP biosynthetic gene cluster is conserved between *Streptomyces* and the plant pathogen *Pseudomonas syringae*; however, L-valine and brassinolide trigger production in strain PG2 but not in *P. syringae*. Together, the inducible production of DHP, and the protection it confers to host plants, underscores a striking combination of evolutionary conservation and ecological specialization in plant-microbe chemical communication.

## RESULTS

### Colonization of *Arabidopsis thaliana* triggers antifungal activity by an endophytic streptomycete

To establish whether bioactive compounds are produced by bacteria upon root colonization, we screened a collection of streptomycetes isolated from the endophytic root compartment of *A. thaliana* for their ability to produce bioactive compounds that inhibit the growth of the fungal pathogen *Rhizoctonia solani*. Bioactivity was assessed in a dual culture assay by growing isolates of the collection on the edge of a petri dish on minimal medium. Five isolates were inoculated per petri dish, equidistant from one another, with *Streptomyces griseus* and *Streptomyces coelicolor* used as controls. After five days, *R. solani* was inoculated in the middle of the plates. Inhibition of fungal growth was measured after a further three days of incubation. Five out of 35 strains inhibited hyphal growth of *R. solani* **(Fig. S1)**. Following these results, sterile seven-day-old *A. thaliana* seedlings were inoculated with *Streptomyces* spores and then moved to sterilized soil when colonization was visible **(Fig. 1a)**. After seven days, *R. solani* plugs were added to the pots just below the soil surface at 0.5 cm from the stem base. *R. solani* infected sterile control plants, leading to plant death within 10 days. One of the isolates, *Streptomyces* sp. PG2, inhibited root infection by the fungus **(Fig. 1b)**. The four other strains that showed bioactivity against *R. solani* during initial *in vitro* screening did not prevent plant infection **(Fig. S2)**, demonstrating the importance of *in vivo* validation. Based on this finding, *Streptomyces* sp. PG2 was selected for further investigation. To visualize the inhibition of fungal root infections in more detail, the experiment was repeated on agar plates. Seedlings inoculated with *Streptomyces sp.* PG2 spores, non-inoculated sterile *A. thaliana* seedlings, or *Streptomyces sp.* PG2 alone were grown on half-strength MS agar plates, with an agar plug of *R. solani* inoculated in the centre of the plate. While plants or *Streptomyces* alone were readily overgrown by the fungal pathogen, a clear zone of hyphal growth inhibition of the fungus was observed around the plants inoculated with *Streptomyces sp.* PG2 **(Fig. 1c)**.

**Figure 1.**
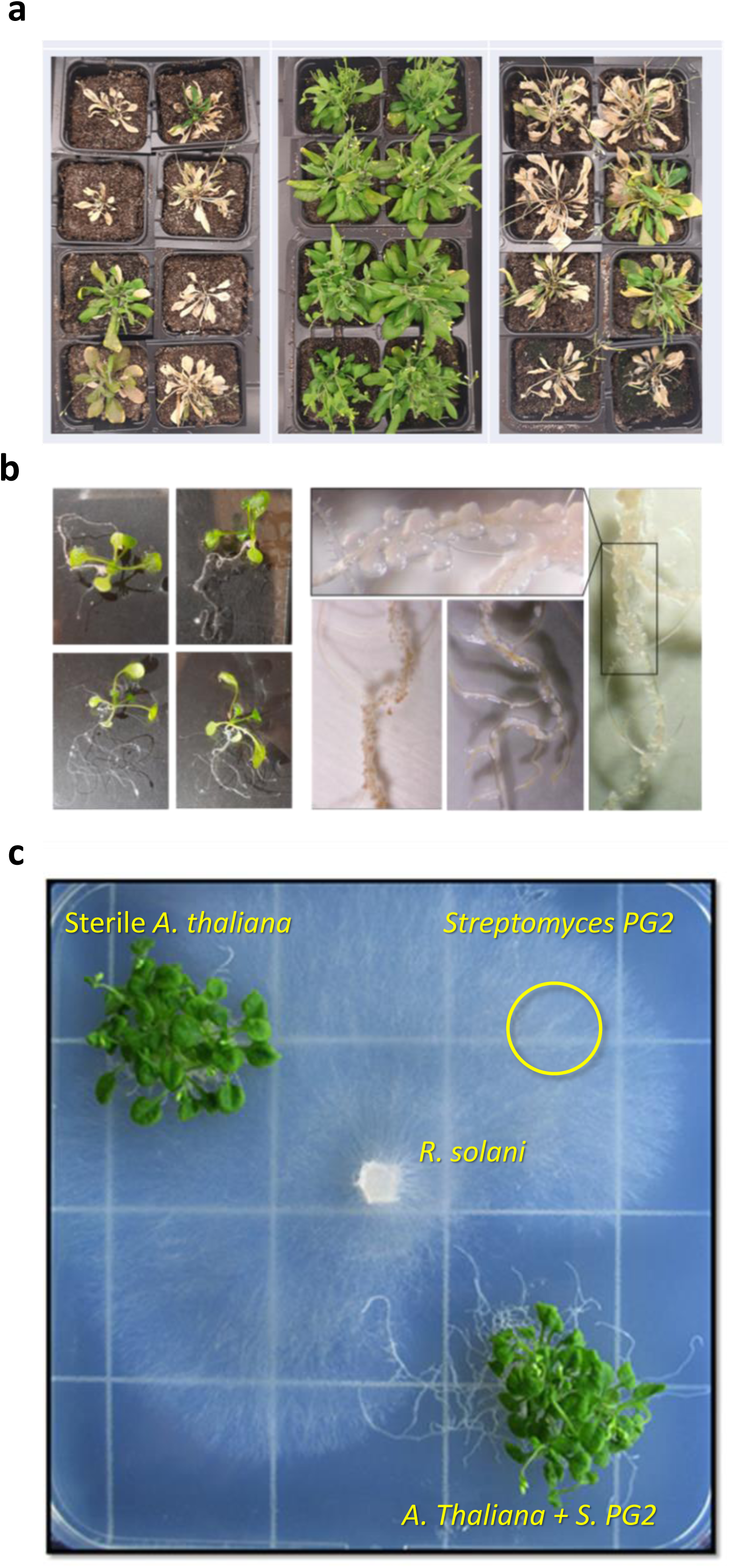
Plant-inducible suppression of *Rhizoctonia solani* infection by *Streptomyces* sp. PG2. **a)** *A. thaliana* seedlings colonized by *Streptomyces* species. The colonization is visible as white spots on the root system. The magnification on the right shows the mycelium growing on the roots. **b)** *In vivo* bioassay in soil. Plants colonized by *Streptomyces* sp. PG2 (middle) are more resistant to infection by the fungal pathogen *R. solani* as compared to sterile plants (left) and those inoculated with a control endophytic *Streptomyces* that does not produce antifungals (right). **c)** Bioassay on agar plate. Top left, *A. thaliana* grown alone; top right, *Streptomyces* PG2 grown alone; bottom right, *A. thaliana* inoculated with *Streptomyces* PG2. *R. solani* was spotted in the middle and allowed to grow for 5 days. Note that while *R. solani* overgrows both the plant and the bacterium alone, it fails to grow near *A. thaliana* colonized by *Streptomyces* sp. PG2.

### Plant colonization elicits the biosynthesis of DHP in *Streptomyces* sp. PG2

We characterized the antifungal activity by investigating the metabolites released by *Streptomyces sp.* PG2 in liquid half-strength MS medium with or without *A. thaliana* seedlings. One-week old spent media were used to assay bioactivity on agar plates. Interestingly, while spent cell-free media from the bacterial cultures partially inhibited fungal growth, no inhibition was seen for the plant-derived media. However, antifungal activity was strongly enhanced in spent media obtained from plant–*Streptomyces* co-cultures **(Fig. 2a)**. These results suggested that the antifungal activity originated from *Streptomyces* sp. PG2, and that its biosynthesis was elicited only in the presence of *A. thaliana*. It is important to note that the plants did not show any signs of distress related to *Streptomyces* sp. PG2 colonization or antimicrobial production, remaining healthy throughout the duration of the experiment, when compared to the control group.

**Figure 2.**
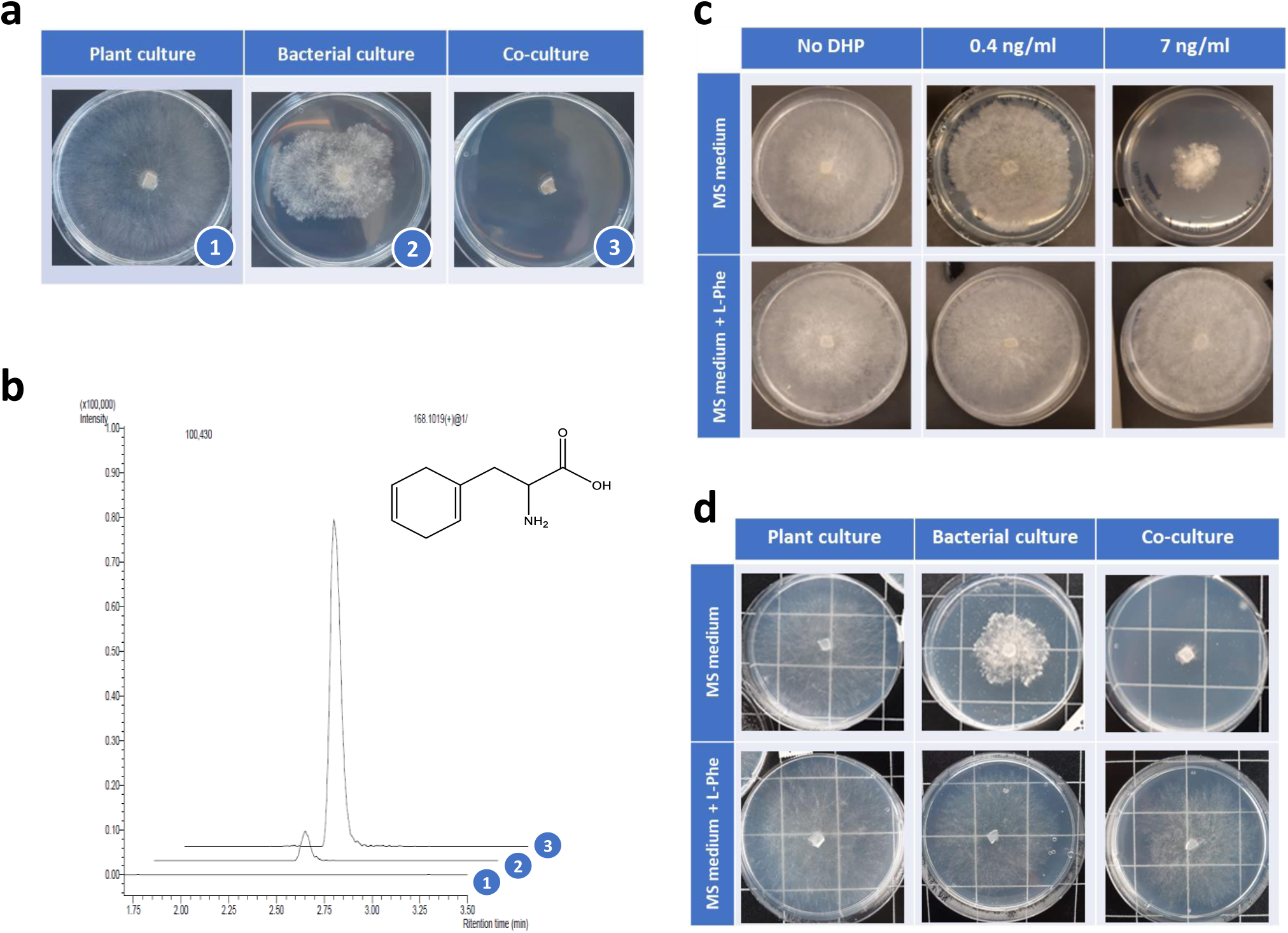
Identification of DHP as an antifungal agent produced by *Streptomyces* sp. PG2. **a)** Bioassays against *R. solani* grown on media with extracts obtained from *A. thaliana* grown alone (left), *Streptomyces* PG2 grown alone (middle), or the plant–*Streptomyces* co-culture (right). Note that *R. solani* fails to grow on agar supplemented with extracts from the plant–*Streptomyces* coculture, while it grows well on extracts from either monoculture. The numbers correspond to the respective LC-MS profile in Panel b. **b)** Extracted ion chromatogram (EIC) of DHP (168.1019 m/z; chemical structure is shown) produced by *Streptomyces* PG2 grown as monoculture (trace 2) or in the presence of *A. thaliana* (trace 3); no peak was detected in samples from pure plant cultures (trace 1). **c)** Antifungal activity of chemically synthesized DHP on growth of *R. solani* and chemical complementation with L-Phe. Top, *R. solani* growth is impaired by increasing concentrations of chemically synthetized DHP, with low inhibition at 0.4 ng/mL and strong inhibition at 7 ng/mL DHP. The control (no DHP) is shown on the left. Bottom, chemical complementation with 2 mM L-Phe fully rescues fungal growth. DHP concentrations are the same as in the top row. **d)** Addition of L-Phe restores growth of *R. solani* in the presence of extracts from liquid-grown cultures. Top row, same as in Panel a, with extracts from *A. thaliana* grown alone (left), *Streptomyces* grown alone (middle) or the co-culture (right). Bottom row, the same as in the top row, but with 2 mM L-Phe. Note that L-Phe again fully restores growth to *R. solani*. These experiments provide strong evidence that DHP is the antifungal agent that inhibits fungal growth.

To elucidate the basis of the antifungal activity, the same extracts were analyzed by LC-MS. Comparison of the metabolomes of the different extracts revealed a significant correlation between a mass feature with an *m/z* of 168.1019 and the level of bioactivity **(Fig. 2a–b)**. This mass feature corresponded to the bioactive compound 2,5-dihydro-L-phenylalanine (DHP)^25^. DHP was not present in extracts obtained from the plant alone and in only small quantities in the *Streptomyces* extracts, while it was strongly enriched in the extracts of the co-culture **(Fig. 2b)**. The antifungal activity of DHP likely arises from false feedback inhibition of the shikimate pathway as it serves as an analogue of the amino acid phenylalanine (Phe)^26^. DHP interferes with translation as it can be incorporated into proteins instead of Phe^27^, which stops fungal growth via disruption of microtubule assembly^28^.

To validate the activity of DHP towards *R. solani*, we used chemically synthesized DHP as a quantitative standard in LC-MS and showed that approximately 0.4 ng/mL (2.4 nM) of DHP was produced in the individual bacterial cultures and approximately 7 ng/mL (42 nM) in the co-culture of the bacteria and *A. thaliana*. Administration of the same concentrations of pure DHP to *R. solani* cultures reproduced the inhibition pattern described in **Fig. 2a**, confirming the sensitivity of the fungus to DHP at the concentrations measured in our samples **(Fig. 2c)**. To further validate that DHP was indeed the sole antifungal agent, and that it acts by mimicking L-Phe, we supplemented the pure DHP samples with 2 mM L-Phe, which fully suppressed the antifungal activity in the extracts, strongly suggesting that L-Phe antagonized DHP and also that no other antifungals were produced, at least not at levels sufficient to inhibit growth of *R. solani* **(Fig. 2c)**. Similarly, L-Phe complementation of samples from pure bacterial/plant culture and bacterial-plant co-culture fully rescued the fungal growth **(Fig. 2d)**, suggesting that DHP was most likely solely responsible for the antifungal activity of the extracts.

### Identification of the DHP gene cluster in *Streptomyces* sp. PG2

DHP production has been reported for *Streptomyces*^25, 29^ and certain Gammaproteobacteria^30–32^. The DHP biosynthetic gene cluster (BGC) was identified for the Gammaproteobacteria *Erwinia amylovora* and *Photorhabdus luminescens*^30^, but not in Actinobacteria. To identify the DHP BGC in *Streptomyces*, we obtained the whole genome sequence of *Streptomyces* sp. PG2, with 7 contigs and a genome size of 10.58 Mb. antiSMASH^33^ predicted 29 BGCs, but no BGC for DHP was detected **(Fig. S3)**. However, searching the PG2 genome with the DHP BGC from *E. amylovora* and *P. luminescens* as queries resulted in the identification of a putative BGC, of which five genes showed significant homology to genes of *E. amylovora* and *P. luminescens*. Of these, *plu3042* and *plu3043* encode a branched-chain amino acid aminotransferase IlvE and a prephenate decarboxylase, respectively, and were previously reported to be necessary for DHP production in *P. luminescens*^30^. The genes are preceded in the operon by the *plu3041* (*paaK*) gene, which encodes phenylacetate-coenzyme A ligase PaaK. The genomic environment of the locus and the conservation of the genes are shown in **Fig. 3a**.

**Figure 3.**
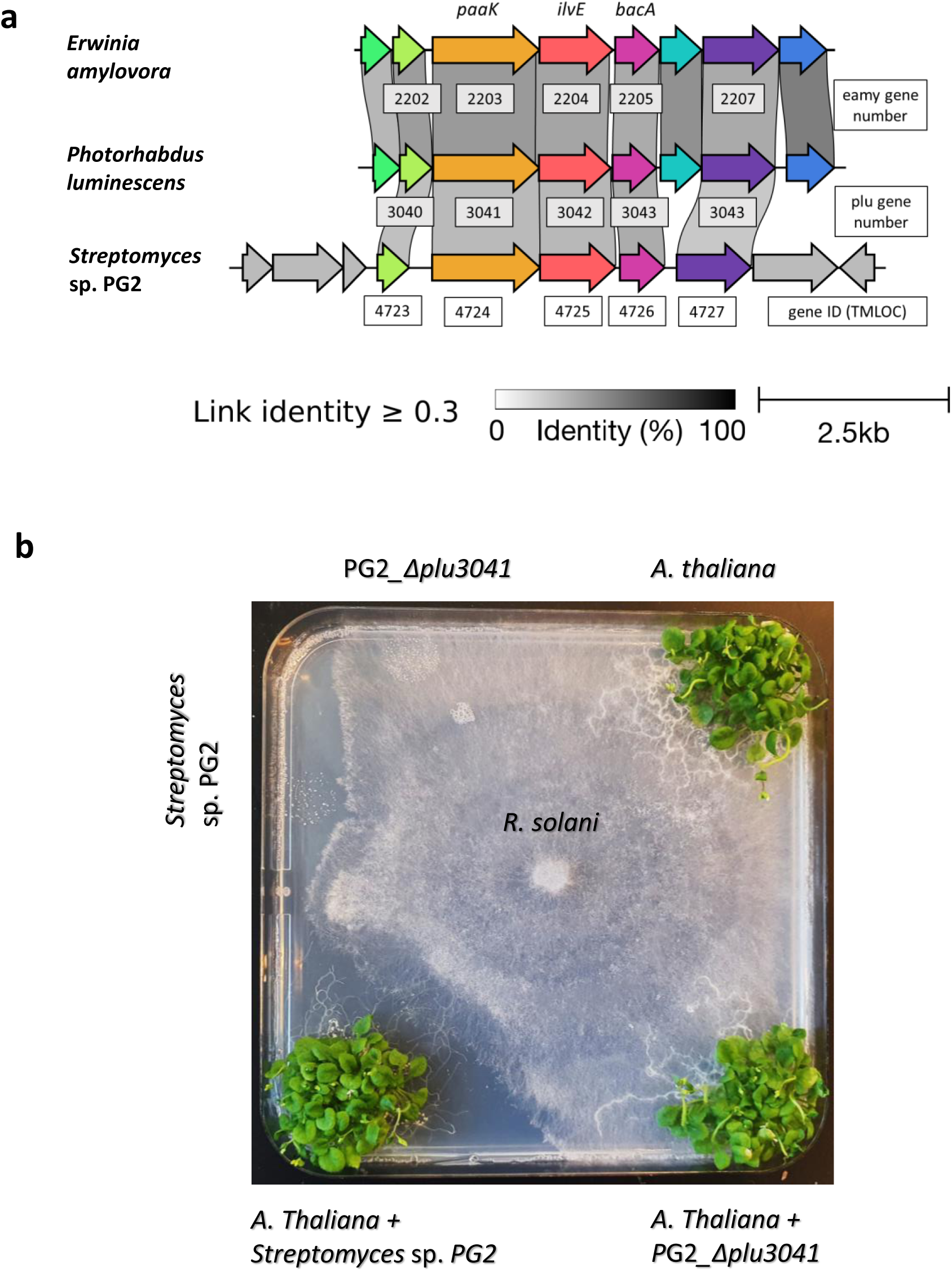
DHP cluster conservation and mutational analysis. **a)** Alignment and conservation of DHP gene clusters identified in *Erwinia amylovora, Photorhabdus luminescens* and *Streptomyces* sp. PG2. The *plu3041-3043* operon is highlighted in blue. **b)** The DHP gene cluster is responsible for the antifungal activity. DHP Knock-out mutant PG2_Δ*plu3041* of *Streptomyces* sp. PG2 fails to inhibit growth of *R. solani*. Within 3 days of growth, the fungus overgrows the plant alone (top right), the mutant alone (top left) and the plant grown in co-culture with the mutant (bottom right). In line with the results shown in Figure 1b, the fungus cannot grow near the plant–*Streptomyces* co-culture due to production of antifungals (bottom left). Slight inhibition of the fungus is shown for *Streptomyces* sp. PG2 grown alone (left).

To ascertain that this gene cluster directs the biosynthesis of DHP, a mutant was created whereby the orthologue of *plu3041* was replaced by double recombination with the apramycin resistance cassette *aac(C)3* using the unstable multi-copy plasmid pWHM3^34^ (see Materials and Methods for details). This resulted in a mutant whereby nt positions -4 to +1326 bp relative to the start of gene TMLOC_04724 were replaced by the apramycin resistance cassette, which is expected to also transcriptionally block the downstream genes *plu3042-3043*. The resulting mutant was designated PG2_Δ*plu3041*. The null mutant was then grown alone or in combination with *A. thaliana* seedlings and challenged with *R. solani*. Mutant PG2_Δ*plu3041* failed to inhibit the growth of the pathogen **(Fig. 3b)**. Metabolomic analysis of PG2_Δ*plu3041* confirmed that the mutant failed to produce any DHP, alone or in combination with *A. thaliana* **(Fig. S4a)**. Finally, to ensure that the phenotype was solely due to the mutation of gene TMLOC_04724 or the disruption of the expression of genes in this area, we genetically complemented the mutant, by introducing the whole gene cluster back into the knock-out mutant. To do so, we created a pSET152^35^ derivative containing a targeted region from TMLOC_04720 to TMLOC_04729, namely pE-DHP_Hyg. The plasmid was then introduced into PG2_Δ*plu3041* via conjugation. Genetic complementation restored DHP production in the mutant, enabling DHP production specifically in response to plant co-cultivation at levels equivalent to those of the wild-type strain **(Fig. S4a** and **S5b)**, showing that indeed the BGC was solely responsible for DHP biosynthesis. As expected, the plant-dependent antifungal activity against *R. solani* was also restored by the genetic complementation of the BGC **(Fig. S6)**. Taken together, these data show that the BGC is required for production of DHP and for inhibition of *R. solani* growth.

### Colonization-dependent elicitation of DHP biosynthesis in other plant-associated bacteria

A search with three core genes of the DHP BGC against the RefSeq database^36^ showed that around 1% (n=720) of all bacterial genomes (including WGS records) contain a likely DHP gene cluster. DHP gene clusters were found in the phyla Actinobacteria, Gammaproteobacteria and Betaproteobacteria. The major genera were *Erwinia* (230) and *Pseudomonas* (186) within the Gammaproteobacteria, *Streptomyces* (94) within the Actinobacteria and *Photorhabdus* (73) within the Betaproteobacteria. The diversity of the gene clusters was analyzed using BiG-SCAPE^37^ **(Fig. 4a)**, with a 0.3 clustering distance cut-off (default value). The DHP gene clusters found in *Streptomyces* genomes showed significant diversity, separating into multiple network clusters, whereby a large number of clusters shared almost the same sequence (distance = 0). Many of the clusters were found in genomes obtained in a few “BioProjects” for comparative genome analyses, especially for the plant pathogen *Pseudomonas syringae*. Of note, most of these projects are collections of bacterial samples from plants **(Table S1)**, suggesting an association between the DHP BGC and plant-associated bacteria. However, further validation using an unbiased database is necessary to ascertain this connection.

**Figure 4.**
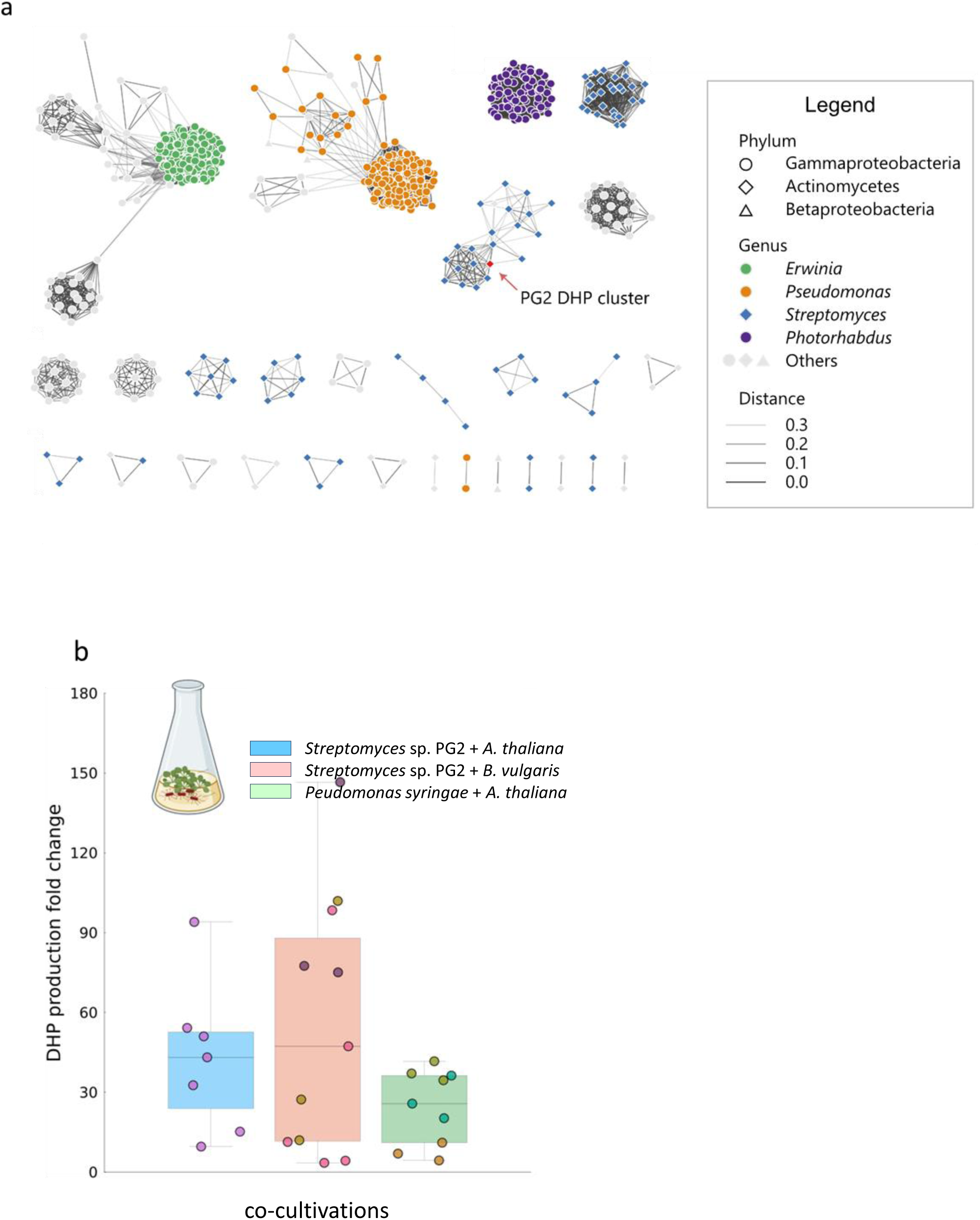
Phylogenomic analysis of DHP gene cluster distribution and plant-mediated elicitation of DHP biosynthesis. **a)** Network of DHP clusters from all bacterial genomes produced by BiG-SCAPE. DHP gene clusters were analyzed with a default cut-off (0.3) (BiG-SCAPE 10.1038/s41589-019-0400-9). Each node represents a single DHP gene cluster from one genome. The phyla are coded using node shapes, genera are coded using node colors. BiG-SCAPE calculated cluster distances are shown as different line darkness. The PG2 genome was added to the collection before analysis. Singletons are not included in the graph. **b)** Fold-change in DHP production in several co-culture conditions compared to pure bacterial cultures. For *Streptomyces* PG2 grown in co-culture with *A. thaliana* (blue box), each data point represents the outcome of an experiment conducted in technical triplicate. For *Streptomyces* sp. PG2 with *B. vulgaris* and *Pseudomonas* FF5 with *A. thaliana* (red and green boxes, respectively), the experiment was run three times, whereby replicates belonging to the same experiment are represented with the same color. DHP quantification was performed via integration of the LC-MS peak.

DHP biosynthesis is conserved in plant-associated phyla such as *Pseudomonas*. We therefore wondered whether plant colonization-dependent production of DHP was also conserved in other phyla. We analysed DHP production in the plant pathogen *Pseudomonas syringae* pv. *syringae* strain FF5. *P. syringae* FF5 harbors the DHP BGC, but so far has never been reported to produce DHP^30^. To see if expression of the DHP BGC of *P. syringae* FF5 could be elicited upon plant colonization, the strain was grown in MS medium in monoculture or in co-culture with *A. thaliana* seedlings. While DHP production by FF5 monocultures was extremely low, near background noise levels, the production of DHP was strongly enhanced in co-culture with the plant, showing up to 41-fold increase as compared to the monoculture **(Fig. 4b)**. Collectively, these findings reveal conservation of elicitation of the DHP gene cluster specifically upon plant colonization across bacteria belonging to different phyla.

### Exploring DHP elicitation by *Arabidopsis thaliana*

We next investigated the cues that could mediate DHP induction upon colonization. To investigate whether interaction with live plants is required, strain PG2 was grown in the presence of either spent media from axenic plants that contain metabolites released by the plant irrespective of colonization, or in the presence of dead plant material (following freezing). In both conditions, DHP production was not induced **(Fig. S5a)**. This excludes the possibility that DHP is elicited either by physical contact with inactive plants, or by plant metabolites produced in the absence of the bacterium. Since plant-microbe interactions often involve volatile compounds (VCs), we next conducted a split-plate assay where *Streptomyces* sp. PG2 was separated from PG2-inoculated *A. thaliana* by a plastic barrier. In this setup, the only way for the plant to activate DHP biosynthesis in the bacterium would be through the air, i.e. by producing VCs. However, *Streptomyces* sp. PG2 exposed to VCs released from PG2-colonized *A. thaliana* did not produce more DHP than the control conditions (sterile *A. thaliana*, *Streptomyces* sp. PG2 monoculture or MS medium without biomass, **Fig. S5a**). These results show that VCs are not the (sole) causative agents of DHP elicitation.

We then sought to investigate whether the elements required for recognition of the plant signals were conserved in the BGC itself. We introduced the plasmid pE-DHP_Hyg carrying the DHP BGC into *Streptomyces coelicolor* M145. This model streptomycete^38^ does not have its own copy of the DHP gene cluster and does not produce DHP. Heterologous expression of the DHP BGC resulted in the production of DHP in *S. coelicolor*, providing further evidence that the BGC is solely responsible for DHP production in *Streptomyces.* Excitingly, DHP levels were significantly enhanced when *S. coelicolor* was co-cultivated with *A. thaliana* **(Fig. S4b** and **S5b)**.

Finally, to explore whether plant colonization-dependent DHP elicitation was specific for *A. thaliana* or could also be mediated by colonization of other plant species, *Streptomyces* sp. PG2 was grown in association with sugar beet (*Beta vulgaris*). Again, DHP production was enhanced to a similar extent as during interaction with *A. thaliana* **(Fig. 4b)**. Collectively, these results strongly suggest that interaction with live plants is critical for DHP induction and that plant colonization-dependent elicitation of the BGC is conserved between streptomycetes, including in species that do not have the DHP BGC.

### DHP is elicited by L-valine and brassinolide hormones in *Streptomyces* sp. PG2

Our results suggested that DHP is elicited by host-specific metabolites produced by live plants. To systematically test this hypothesis, we employed high-throughput elicitor screening (HiTES), a chemical genetics approach that evaluates how individual molecules perturb secondary metabolism^39^. Building on recent results with ecology-guided HiTES (Eco-HiTES), which has revealed chemical communication mechanisms between marine bacteria and algae using host-derived metabolites as elicitors^40^, we created a custom plant elicitor library comprising 103 common plant metabolites **(Fig. 5a**, **Table S2)**. Screening this library against *Streptomyces* sp. PG2 revealed five compounds that strongly induced DHP production in the primary assay: the plant hormone brassinolide, the branched-chain amino acid (BCAA) L-valine, vanillic acid, and two plant metabolites not naturally produced by *A. thaliana*—linamarin and pterostilbene **(Fig. 5b–c)**. We focused further on the three *A. thaliana* metabolites and, while vanillic acid did not reproducibly elicit DHP, both brassinolide and L-valine triggered DHP production in a dose-dependent manner, independent of bacterial growth effects **(Fig. 5d–e, Fig. S7a–b)**. Related compounds, including epibrassinolide and the BCAA L-isoleucine, also elicited DHP production, albeit at lower levels **(Fig. S7c–d)**. While L-leucine elicited a slight increase in DHP, the response did not reach statistical significance **(Fig. S7d)**.

**Figure 5.**
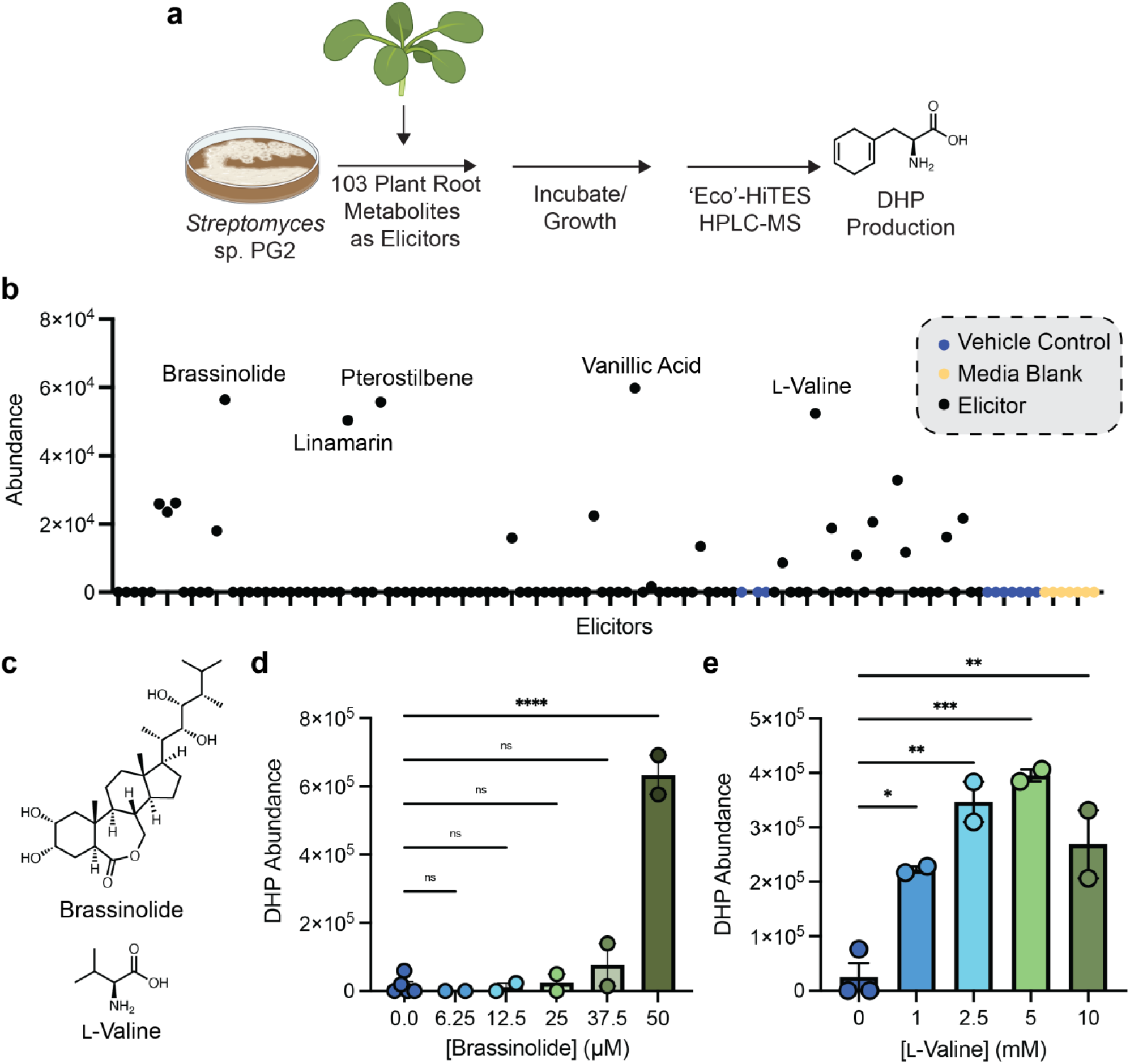
Eco-HiTES identifies L-valine and brassinolide as elicitors of DHP in *Streptomyces* sp. PG2. **a)** Overview of the Eco-HiTES approach with *Streptomyces* sp. PG2 and the plant elicitor library. **b)** Abundance of DHP in each elicitor, control, or media condition tested in the Eco-HiTES screen. **c)** Chemical structures of brassinolide and L-valine. **d–e)** Dose response curves showing how DHP abundance increases with increasing concentrations of **d)** brassinolide and **e)** L-valine. For **d–e**, samples were grown as biological triplicates or duplicates; error bars represent the SE of the means. Significance for **d–e** was determined by Dunnett’s multiple comparison test and for each, \*\*\*\**P* ≤ 0.0001, \*\*\**P* ≤ 0.001, \*\**P* ≤ 0.01, \**P* ≤ 0.05, & ns = not significant.

We then tested the effect of the elicitors on the antifungal activity of PG2 in plate assays, using a similar setup as shown in Fig. 2. *Streptomyces sp.* PG2 was grown in liquid half-strength MS medium containing either 5 mM L-valine, 50 μM brassinolide or no added elicitor. One-week-old spent media were used to assay bioactivity on agar plates against *R. solani*. While brassinolide did not result in a significant change in antifungal production, addition of L-valine led to inhibition of fungal growth **(Fig. S8a)**. Finally, we compared brassinolide mutant plant col-0 06875 and wild-type *A. thaliana* for their ability to stimulate DHP production in PG2. Importantly, while both plants induced DHP production, the level of induction by the brassinolide mutant was significantly reduced **(Fig. S8b)**. Together, these data show that brassinolide and L-valine act as elicitors of DHP production in *Streptomyces* sp. PG2.

To determine whether this response was conserved, we tested *S. coelicolor* expressing the DHP gene cluster. L-valine induced DHP in both PG2 and *S. coelicolor* at comparable levels, but brassinolide did not, suggesting that PG2 possesses additional regulatory pathways absent in *S. coelicolor* **(Fig. S7e–f)**. Finally, we tested whether L-valine or brassinolides elicited DHP in the plant pathogen *P. syringae*, but did not observe any DHP beyond background levels **(Fig. S7g)**, suggesting that *P. syringae* has a disparate regulatory mechanism for inducing DHP biosynthesis. These results suggest that brassinolide elicitation is unique to PG2, whereas L-valine regulation is conserved across the tested streptomycetes, and that *P. syringae* employs a distinct, yet unknown, mechanism for DHP induction.

## DISCUSSION

Members of the genus *Streptomyces*, the largest actinobacterial genus, are well-known producers of a broad range of bioactive metabolites^11, 41^. Given the substantial antimicrobial potential of streptomycetes and their ability to colonize diverse plant species^9, 10, 42–44^, understanding how to harness plant-microbe interactions for crop biocontrol is crucial. Still, despite the exponential growth of microbial genome sequence information, insights into the influence of host-microbe interactions on activating the desired genomic traits in the microbiome remain limited.

In this work, we show that plants stimulate the production of antimicrobials by members of their microbiome, resulting in protection against microbial pathogens **(Fig. 6)**. We identified *Streptomyces* sp. PG2 as an efficient bioprotectant, producing the antifungal DHP specifically in response to plant colonization. Furthermore, introduction of the DHP BGC into *S. coelicolor* that lacks the BGC, led to plant-inducible DHP production, showing conservation of (i) detection of external cues; (ii) subsequent signal transduction; and (iii) regulation of DHP biosynthesis. Elicitation of DHP production was strongly induced by interaction with live plants, but could not be mediated by volatile compounds produced by the plant alone or by *Streptomyces*-colonized plants. We were able to pinpoint two classes of soluble plant metabolites, brassinolide hormones and BCAAs, as potent cues to elicit DHP in *Streptomyces* sp. PG2. Extracts from axenic plants failed to elicit DHP production, likely due to the fact that the concentrations of the elicitors were too low for application in liquid-grown cultures or plate assays.

**Figure 6.**
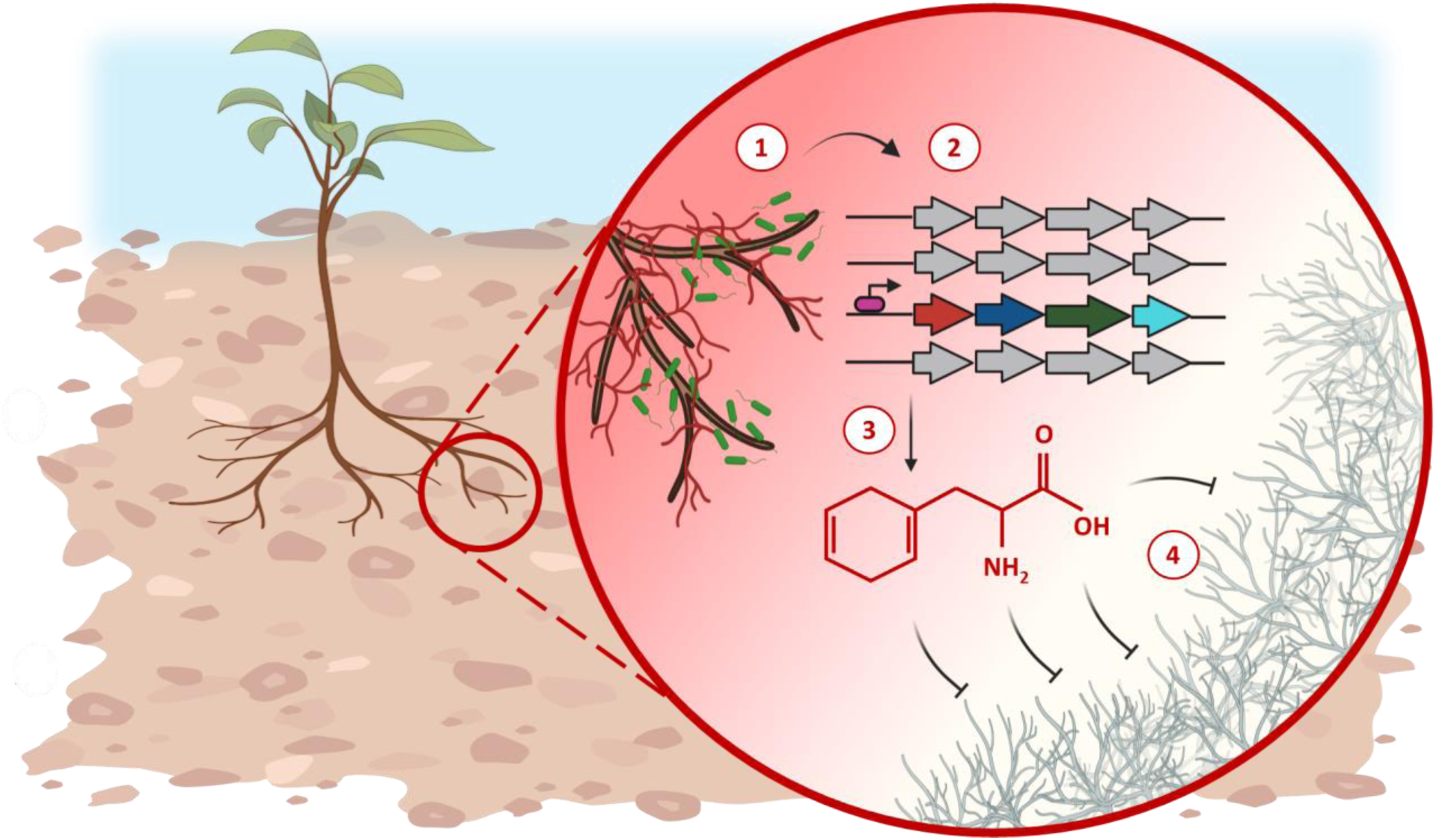
Model for DHP gene cluster activation. (1) Root colonization; (2) the molecular cross-talk between the plant and its associated microbes results in the activation of the DHP gene cluster; (3) DHP is released into the surroundings of the colonized host, and it prevents soil borne pathogens to reach the root system (4).

The elicitation by L-valine may reflect multiple, non-mutually exclusive mechanisms. Metabolically, L-valine could serve as an amino donor in the biosynthesis of DHP. During DHP biosynthesis, the aromatic amino-acid precursor prephenate is converted by a non-aromatizing prephenate decarboxylase to a reduced cyclohexadienyl keto-acid (endocyclic dihydro-hydroxyphenylpyruvate), which is then converted into DHP via a transaminase. BCAAs are amino donors for branched-chain aminotransferases (BCAT-type enzymes), which transfer the amino group from the amino acid to a keto-acid acceptor^45^. Supplying L-valine can thus directly feed the DHP pathway by providing the amino group for the final aminotransferase step, consistent with the known requirement for an amino-acid cosubstrate in the transamination step in H₂Phe biosynthesis^45^.

In parallel, L-valine may function as a regulatory cue that increases BGC transcription. Consistent with a conserved regulatory response, L-valine induced DHP in both PG2 and *S. coelicolor* expressing the DHP cluster, whereas brassinolide did not, suggesting additional PG2-specific regulation. Notably, the *S. coelicolor* cluster encodes TMLOC_04729, a putative leucine-responsive transcriptional regulator (Lrp), raising the possibility that an Lrp-family regulator responds to L-valine (or other BCAAs) to promote DHP biosynthesis. The *P. syringae* DHP BGC does not contain a homolog for this Lrp-family regulator, further aligning with this hypothesis.

Importantly, we discovered that plant colonization-dependent induction of DHP spans streptomycetes and includes the plant pathogen *P. syringae*, illustrating this phenotype is conserved between distant bacterial phyla. Yet, the molecular mechanism between the two is divergent, a difference that can likely be attributed to the fact that *Streptomyces* sp. PG2 is an endosymbiont, while *P. syringae* is a pathogen. Most of the experiments were done with *A. thaliana*. However, we also observed DHP stimulation upon colonization of other plant species, such as *B. vulgaris*, possibly opening the door to broader applications in crop protection.

In line with the notion that DHP serves as a plant-protective agent, phylogenomic analysis revealed an intriguing correlation between the presence of the DHP cluster and a plant-associated lifestyle. We could only identify the BGC in the genomes of species of *Erwinia* (230), *Pseudomonas* (186), *Streptomyces* (94) and *Photorhabdus* (73), many of which are known to be plant-associated^11, 17, 46, 47^. Most of the DHP clusters from *Pseudomonas* were found in the genomes of bacteria obtained from plant samples. *Erwinia* is a genus of bacteria mostly composed of plant pathogenic species, whereas for *Streptomyces* it is hard to draw a conclusion as the details underlying the ecology of many strains are unknown. *Photorhabdus* — best known as a symbiont of nematodes — was recently shown to respond to plant signals, contributing to chitin degradation and inhibition of fungal growth^48^. *Photorhabdus* species indirectly interact with plant signals through their symbiotic relationship with obligate parasites of herbivorous insect larvae. As a result, *Photorhabdus* and its symbiont *Heterorhabditis* are being studied as potential biocontrol agents^46, 49^ for crop protection. We propose that both beneficial and pathogenic microbes may exploit DHP production to diversify their metabolic capabilities, thereby capitalizing on plant cues to enhance the resilience of their host or to improve their competitiveness within the rhizosphere, respectively.

Heterologous expression of the gene cluster in the DHP non-producer *S. coelicolor* showed that biosynthesis was activated in a plant-dependent manner suggesting that at least part of the regulatory circuitry is located within the BGC itself. We speculate that the Lrp-family regulator is the local regulator within the BGC. Regulatory networks coordinating critical cellular processes are typically preserved over evolutionary time scales and are often regulated by ancestral transcription factors^55^. The evolutionary conservation of the control of its biosynthesis indicates that DHP may play an important role in the lifestyle of plant-associated microbes. We recently showed that regulatory networks may be harnessed for the functional prediction of BGCs, using iron control as the basis for the discovery of novel BGCs involved in siderophore biosynthesis^50^. Therefore, elucidating how the plant-dependent activation of antifungal biosynthesis is governed will enable the search for BGCs associated with similar regulatory networks as candidates involved in plant protection.

To the best of our knowledge, our work provides the first example of a natural product that is produced by microbes specifically in response to plant colonization, and whose stimulation is conserved both in beneficial and pathogenic plant-associated microbes. By leveraging Eco-HiTES, we have demonstrated the power of ecology-guided discovery to uncover regulatory pathways of cryptic metabolites. Through this approach, we uncovered how the DHP biosynthetic pathway is activated in the endosymbiotic *Streptomyces* sp. PG2, whereas the plant pathogen *P. syringae* appears to rely on a different regulatory mechanism. This highlights unresolved questions regarding the specificity of chemical signals that hosts use to communicate with either symbionts or pathogens. Deeper investigation into the chemical interplay regulating this phenomenon will be required to fully elucidate this inter-kingdom dialogue. Our study offers important new insights into the dynamics occurring amongst several players that populate the rhizosphere and motivates further exploration of the biosynthesis of cryptic bioactive molecules specifically during plant-microbe interactions. Our findings also offer great promise for novel approaches for the sustainable protection of crops against fungal pathogens and pests.

## MATERIALS & METHODS

### Microbial strains and culturing conditions

The isolation of endophytic bacteria was employed as described previously ^51^. Briefly, root tissue of *A. thaliana* ecotype mossel (Msl) obtained from a natural ecosystem was cleaned, sonicated and ground with mortar and pestle in 1 mL phosphate buffer (per liter: 6.33 g of NaH_2_PO_4_·H_2_O, 10.96 g of Na_2_HPO_4_·2H_2_O and 200 μL Silwet L-77). The plant material was spread onto the surface of a range of selective isolation media. Initial selection was done on media supplemented with the antifungal agent nystatin (50 µg/mL) and the antibacterial agent nalidixic acid (10 µg/mL). Plates were incubated at 30 °C for 4–25 days. Single strains were isolated based on morphology, grown on soya flour mannitol (SFM) for seven days at 30 °C, and spores were collected^52^. The 35 strains that were obtained were grouped under the name ATMOS collection. *Streptomyces coelicolor* A3(2) M145^38^ was obtained from the John Innes Centre strain collection and the host for heterologous expression experiments. The fungal plant pathogen *Rhizoctonia solani* was used in this work as an antifungal activity reporter. The fungus *R. solani* AG2-2IIIB was obtained from the collection of the Netherlands Institute of Ecology (NIOO-KNAW). Agar plugs with fungal hyphae used as inoculum in the bioassays were prepared by growing *R. solani* on 1/5th potato dextrose agar (PDA) medium for three days at 25 °C. *Pseudomonas syringae pv. syringae* strain FF5 was obtained from the IHSM-UMA-CSIC, Malaga, Spain.

### Plant materials and growth conditions

Wild-type *Arabidopsis thaliana* Columbia-0 (Col-0) ecotype and *Beta vulgaris* (sugar beet) cultivar Camelia were employed in this study. *A. thaliana* seeds were obtained from Netherlands Institute of Ecology (NIOO-KNAW). Sugar beet seeds were obtained from the Sugar beet Research Institute (IRS, The Netherlands), and were kindly provided by Victor Carrion Bravo group. The seeds were surface-sterilized by soaking for 10 min in 40 % bleach. After bleach removal, they were rinsed five to seven times with sterile distilled water, and stored at 4 °C. Half-strength Murashige-Skoog (MS) medium was used for plant growth. One liter of MS medium contains: 2.2 g MS powder (Duchefa), 10 g sucrose; the pH was adjusted to 5.7 and 10.8 g Micro Agar (Duchefa) was added for solid medium prior to autoclaving. Following the seed sterilization, all the plants in this work were germinated on half-strength MS plates for one week. After germination, seedlings for LC-MS analysis and bioassays were moved to liquid cultures in Erlenmeyer flasks and incubated in shaking mode. For soil experiments, one-week-old seedlings were transferred from plates into liquid medium to 24 multi-well plates, incubated in shaking mode for one week and, eventually, moved on twice-autoclaved mixture of 9:1 substrate soil and sand (Holland Potgrond). *A. thaliana* and *B. vulgaris* on agar plates, liquid medium or soil, were cultivated at 21 °C, 16 h photoperiod, and 50% relative humidity; *B. vulgaris* was grown at 24 °C, a 16 h photoperiod, and 70% relative humidity.

### Bioassays

All bioassays are presented in the **Supplemental Methods**. These include ***in vitro* bioassays**, namely: ATMOS collection screening; co-culture spent media; DHP bioactivity assays; and chemical complementation with L-Phe to chemically complement DHP and rescue the fungal growth; and ***in vivo* bioassays**, namely agar plate assays and in-soil assays.

### Analysis of secondary metabolites

Pure bacterial/plant cultures and plant-bacteria co-cultures were set up to analyze the differential metabolite production underlying the increased bioactivity of plant-bacterial co-culture spent media. Samples were prepared in triplicate. In brief, 15 one-week-old seedlings were moved to flasks in 15 ml half-strength MS liquid medium and inoculated with 4.5 x 10^3^ *Streptomyces* spores. For plant-*Pseudomonas* FF5 co-cultures, 50 µl of FF5 overnight culture adjusted to OD_600_ 0.5 in half-strength MS medium was added to the medium instead of *Streptomyces* spores. Pure bacterial cultures were prepared by adding the same amount of inoculum to sterile media. The flasks were incubated in shaking mode at 21 °C, 16 h photoperiod, and 50% relative humidity (*A. thaliana*), or at 24 °C, 16 h photoperiod, and 70% relative humidity (*B. vulgaris*) for seven days. The experimental setup for the study of possible interaction via the production of **Volatile Compounds** (VCs) was as follows, using the same media. Spores from *Streptomyces* sp. PG2 were inoculated on one side of a Petri dish and *A. thaliana* inoculated with PG2 was grown on the other side, with an impermeable sector dividing them in the middle. After 10 days of incubation, metabolites were extracted from the side with the *Streptomyces* sp. PG2 monoculture and checked for DHP.

For the study of soluble metabolites in liquid-grown cultures, supernatants were collected, 5% w/v Diaion® HP20 (Resindion) was added to the cultures and shaken for three h at room temperature. HP20 was filtered off the liquid media with glass wool, washed with distilled water, then soaked with 100% MeOH three times overnight. For extraction of either soluble or volatile metabolites from solid-grown cultures, the agar was chopped into very small pieces and submerged in 100% methanol three times overnight. MeOH extracts were dried and dissolved in 75% MeOH in H_2_O to a final concentration of 1 mg/mL and filtered (0.22 µm) for LC-MS analysis. Samples from sterile half-strength MS medium were similarly prepared and extracted to serve as negative controls.

LC-MS/MS acquisition was performed as described ^24, 53^. A Shimadzu Nexera X2 UHPLC system was used, with attached PDA, coupled to a Shimadzu 9030 QTOF mass spectrometer, equipped with a standard ESI source unit, in which a calibrant delivery system (CDS) is installed. For details we refer to the Supplemental Methods.

### Construction of the DHP knock-out mutant

The construct for gene disruption was obtained using a method published previously ^54^. In brief, approximately 1.5 kb regions up- and downstream of the gene TMLOC_04724 from *Streptomyces* sp. PG2 genomic DNA were amplified, using primer pairs 1+2 and 3+4, respectively (Table S3). The apramycin resistance cassette *aac(3)IV* flanked by *loxP* sites, was amplified with primer pair 5+6 from pWHM3-*oriT*^34^. The three DNA fragments were assembled into EcoRI-HindIII digested pWHM3 using NEBuilder® HiFi DNA Assembly. The obtained construct was verified by DNA sequencing, transformed into *E. coli* ET12567/pUZ8002 ^55, 56^ and subsequently introduced into in *Streptomyces* sp. PG2 by conjugation ^52^. After double recombination, nt positions 5308194 to 5309523 of the PG2 chromosome were replaced by the apramycin resistance cassette, corresponding to nt positions -4 to +1326 bp relative to the start of gene TMLOC_04724. The correct mutant was selected by resistance to apramycin (50 µg/mL) and named PG2_Δ*plu3041*.

### Expression of the DHP gene cluster in *Streptomyces*

For genetic complementation we cloned the genomic region between TMLOC_04720 - TMLOC_04729 into the conjugative vector pSET152^57^, which harbours an apramycin resistance cassette *aac(C)IV* as selection marker. For this, three contiguous fragments of this region were amplified with primer pairs 7+8, 9+10 and 11+12 (Table S3). The PCR products were assembled using NEBuilder® HiFi DNA Assembly into BamHI-EcoRI-digested pSET152. The plasmid was named pE-DHP and introduced via conjugation into *S. coelicolor* M145. Positive clones were selected on apramycin (50 µg/mL). The construct sequence was confirmed by restriction digestion and Sanger sequencing, and the obtained strain was named SCO-DHP. SCO-DHP was grown in liquid culture with *A. thaliana* seedlings as described for PG2 strain. Pure bacterial/plant cultures and plant-bacteria co-cultures were set up in triplicate as already described for the *in vitro* bioassays, followed by metabolite extraction and LC-MS run.

To ensure the specificity of the mutation in TMLOC_04724 in PG2_Δ*plu3041*, we reintroduced the entire BGC into the mutant. Since the mutant was already resistant to apramycin, a modified version of pE-DHP was created whereby the hygromycin resistance was inserted into the EcoRV site located inside the apramycin resistance gene *aac(C)IV*. The resulting plasmid pE-DHP_Hyg was then introduced via conjugation into PG2_Δ*plu3041*. The ability of the clone to restore DHP production to PG2_Δ*plu3041* was tested by growing the strain in liquid culture with *A. thaliana* seedlings as described for PG2 strain, followed by LC-MS analysis of the culture extracts.

### Computational methods

#### Phylogenomic distribution of the DHP gene cluster

Genomes were analyzed using antiSMASH (version 6.0.1)^33^ to obtain BGC predictions. Program cblaster (version 1.3.18, 10.1093/bioadv/vbab016) was used for searching the DHP cluster against RefSeq database. Default settings of cblaster were used for BLAST hit. In general, for a protein to match query sequences, minimal identity is set to 30%, minimal coverage is set to 50%, e-value should not be higher than 0.01. To reduce the complexity of searches, query term “txid2[orgn]” is used to restrict the search within Bacteria superkingdom. Predictions were used as input for BiG-SCAPE ^58^ (version 1.1.15, 10.1038/s41589-019-0400-9) with default clustering cutoff 0.3, for the creation of a sequence similarity network, with distance matrix cutoff set to 0.25. The resulting full network was visualized by Cytoscape (version 3.10.1) with default layout^59^.

### Eco-HiTES with *Streptomyces* sp. PG2

To prepare the Eco-HiTES screening plates, a single colony of *Streptomyces* sp. PG2 was inoculated in 5 mL of GYM medium (glucose 4 g L^-1^, yeast extract 4 g L^-1^, malt extract 10 g L^-1^) and grown at 30 °C/250 rpm for 3 days. This culture was used to inoculate Murashige-Skoog medium supplemented with 5% tryptic soy broth (MS-TSB) at a starting cell density of ∼0.01. The culture was distributed into two sterile round-bottom, deep 96-well plates in aliquots of 495 µL. To each well, 5 µL of the elicitor or vehicle control (either dimethyl sulfoxide (DMSO), dimethylformamide (DMF), water, or 50:50 water:acetonitrile (ACN)) was added. Secondary metabolite elicitors were screened at a final concentration of 20 µM, while primary metabolite elicitors were screened at 200 µM (Table S2). The culture plates were covered with Breath-Easy sealing membranes (Sigma) and incubated at 30 °C/250 rpm for 6 days. After incubation, the plates were centrifuged at 2,000 x *g* for 20 minutes, the supernatants were transferred to fresh 96 well plates, diluted with MeOH (7% final), filtered over a 96-well filter plate (0.25 µm, PTFE, Pall corporation), and the filtrate was collected in a 96-well plate. The 96-well plates were covered with 96-well cap mats (Thermo) and 4 µL was injected onto a high resolution Agilent 6540 Accurate Mass QTOF system, consisting of an automated liquid sampler, a 1260 Infinity Series HPLC system, a diode array detector, a JetStream ESI source, and a 6540 Series QTOF, equipped with an analytical Hypercarb™ Porous Graphitic Carbon column (100 x 4.6 mm, 3 µm) with a flow rate of 0.7 mL min^-1^ and an initial elution gradient from 95:5–60:40 water:ACN over 5 minutes, followed by a gradient from 60:40–100:0 water:ACN over the next 2 minutes, with a final isocratic step of 0:100 water:ACN for 3 minutes. All solvents contained 0.1% formic acid. HR-MS data were acquired in positive ion mode with a scan range of 100–3000 *m*/*z*. DHP abundance was quantified by the area under the curve (AUC) of extracted ion chromatograms (EICs), which were generated from each total ion chromatograms (TICs) using the summation of the [M + H]^+^ and [M + Na]^+^ (168.102 and 190.084, respectively) at a retention time of ∼4.2 minutes with a scan width of ± 20 ppm using Agilent MassHunter Workstation. The retention time of DHP was validated by examining supernatants of *Streptomyces* sp. PG2 WT versus PG2_Δ*plu3041*.

### Validation and dose-response of DHP elicitors in *Streptomyces* sp. PG2, *S. coelicolor, and P. syringae*

To validate DHP elicitors, cultures of *Streptomyces* sp. PG2 were grown identically to those described in the ‘Eco-HiTES screen with *Streptomyces* sp. PG2’ section, with the following modifications. Varying concentrations of elicitors were tested in duplicate or triplicate. Brassinolide was tested at 50, 37.5, 25, 12.5, and 6.25 µM with 1% DMSO as a vehicle control; epibrassinolide at 62.5, 50, 37.5, and 25 µM with 1% DMSO as a vehicle control; while L-valine, L-isoleucine, and L-leucine were tested at 10, 5, 2.5, and 1 mM with water as the vehicle control. After 6 days of growth, supernatants were collected, prepared, and analyzed via HPLC-MS as described above. The same protocol was followed for testing *S. coelicolor* harboring the DHP BGC with brassinolide and L-valine. To test whether brassinolide or L-valine modulated growth significantly in *Streptomyces* sp. PG2 or *S. coelicolor* harboring the DHP BGC, cultures were set up in the same manner, and their wet cell pellet weight was measured at day 6. To test whether brassinolide or L-valine elicited production of DHP in *P. syringae* FF5, a single colony was inoculated in 5 mL of LB medium and grown at 30 °C/200 rpm for 18 hours. This culture was used to inoculate MS-TSB at a starting cell density of ∼0.01, which was distributed into sterile, round-bottom, deep 96-well plates in aliquots of 495 µL, and 5 µL of the relevant elicitors were added. Brassinolide was tested at 50, 25, 12.5, and 6.25 µM with 1% DMSO as a vehicle control, and L-valine was tested at 10, 5, 2.5, and 1 mM with water as the vehicle control, in duplicate. The cultures were grown at 30 °C/200 rpm for 18 hours, and the supernatants were prepared and analyzed by HPLC-MS in the same manner as described in the ‘Eco-HiTES screen with *Streptomyces* sp. PG2’ section.

## Supporting information

All supplemental figures and tables

## ACKNOWLEDGEMENTS

We are grateful to Victor Carrion Bravo for providing *Pseudomonas syringae* and sugar beet seedlings. The work was supported by grant OCENW.GROOT.2019.063 from the Netherlands Organization for Scientific Research (NWO) to JMR and GPvW as well as the US National Institutes of Health (grant R35 GM152049 to MRS).

