## Supplementary material for "Antifungal biosynthesis by root-associated *Streptomyces* and *Pseudomonas* is elicited upon plant colonization": All supplemental figures and tables

Belonging to the manuscript

**SUPPLEMENTAL METHODS**

**Bioassays**

***In vitro bioassays***

ATMOS collection screening. To assay antifungal production by isolates of the ATMOS collection, 5 µL of spore stock (1 x $10^{10}$ cfu/mL) were inoculated at the edge of minimal media[1] agar plates, five strains per plate. Plates were incubated for five days at 30 °C, and subsequently *Rhizoctonia solani* was inoculated in the middle of each plate. Inhibition was calculated after three days of growth by measuring the distance in millimeters between the edge of the *Streptomyces* colony and fungal mycelium.

Co-culture spent media. Pure bacterial cultures and plant-bacteria co-cultures were set up to compare the bioactivity of the spent media. *A. thaliana* seedlings were germinated on MS plates. After seven days the germinated seedlings were transferred to liquid cultures in 100 mL Erlenmeyer flasks containing 15 mL half-strength MS liquid medium, 15 seedlings and 4.5 x $10^3$ cfu *Streptomyces*. Plant monocultures were prepared in the same way but without bacteria. Bacterial monocultures were prepared by adding the same inoculum to sterile media. Experiments were performed in triplicate. The flasks were incubated while shaking at 21 °C, 16 h photoperiod, and 50% relative humidity. After seven days the supernatants were collected, filtered (0.22 µm) and supplemented with 10 ml sterile agar solution (15 g/l). The mixtures were poured into Petri dishes. A control plate was poured by mixing half-strength MS medium with the agar solution. An agar plug from a three-day-old *R. solani* culture was inoculated in the center of the plates. The plates were incubated at 28 °C, and growth inhibition measured three days later, when the fungus fully covered the control plates. Images were analyzed with Fiji [2]. Fungal growth inhibition was estimated as follows: $I = (C-R)/C \times 100$, where I is the pathogen inhibition percentage, C is the area of the plate covered by the fungal mycelium in the control plate, R is the growth of *R. solani* measured in the test plates.

DHP bioactivity. This bioassay was conducted to estimate *R. solani* resistance to different concentrations of DHP. Here, the plant pathogen was cultured on 15 g/L agar, supplemented with 0.2, 0.4, 2, 7 ng/mL DHP. *R. solani* growth on agar plates was used as control.

Chemical complementation. In order to assay the ability of L-Phenylalanine to chemically complement DHP and rescue the fungal growth, the plates were supplemented with 2 mM L-Phe before the inoculation of the fungus.

***In vivo bioassays***

Agar plates. Two groups of 15 one-week-old seedlings were transferred at the opposite corners of a square plate, on half-strength MS liquid medium. One of the two was inoculated with 4.5 x $10^3$ *Streptomyces* spores in 100 µL of sterile demi water, while the control was treated with sterile demi-water. After seven days of incubation, *R. solani* was inoculated in the centre of the square agar plate. Plates were incubated at 21 °C, with a 16 h photoperiod and 50% relative humidity to enable regular plant growth. The fungal growth inhibition halo was visible within five days.

In soil. Sterile seeds were germinated on half-strength MS plates and moved to 24 multi-well plates, one seedling per well, in half-strength MS liquid medium. In this step, the seedlings were inoculated with the streptomycete under investigation. Every seedling was inoculated with 3 x $10^3$ spores, ensuring equal distribution. Control seedlings followed the same procedure, omitting the spores inoculation. After one-week-incubation in shaking mode the seedlings were visually analyzed, and the ones showing a visible bacterial colonization on the roots were transferred to sterile soil. The same number of sterile seedlings were moved to soil as control. Ten days later, one plug of *R. solani* was inserted in the soil, close to the stem of the plants. Within one month, the survival ratio of the inoculated plants was calculated.

**LC-MS analysis**

LC-MS/MS acquisition was performed as described [3, 4]. A Shimadzu Nexera X2 UHPLC system was used with attached PDA, coupled to a Shimadzu 9030 QTOF mass spectrometer, equipped with a standard

ESI source unit. Extracts were dissolved in MeOH to a final concentration of 1 mg/mL, and 2 µL were injected into a Waters Acquity HSS $C_{18}$ column (1.8 µm, 100 Å, 2.1 × 100 mm). The column was maintained at 30 °C, and run at a flow rate of 0.5 mL/min, using 0.1% formic acid in $H_2O$ as solvent A, and 0.1% formic acid in acetonitrile as solvent B. A gradient was employed for chromatographic separation starting at 5% B for 1 min, then 5–85% B for 9 min, 85–100% B for 1 min, and finally held at 100% B for 3 min. The column was re-equilibrated to 5% B for 3 min before the next run was started. The LC flow was switched to the waste for the first 0.5 min, then to the MS for 13.5 min, then back to the waste to the end of the run. The PDA acquisition was performed in the range 200–600 nm, at 4.2 Hz, with 1.2 nm slit width. The flow cell was maintained at 40 °C.

The MS system was tuned using standard NaI solution (Shimadzu). The same solution was used to calibrate the system before starting. Additionally, a calibrant solution made from ESI tuning mix (Sigma-Aldrich) was introduced through the CDS system, the first 0.5 min of each run, and the masses detected were used for post-run mass correction for the file, ensuring stable accurate mass measurements. System suitability was checked by including a standard sample made of 5 µg/mL paracetamol, reserpine, and sodium dodecyl sulfate; which was analyzed regularly in between the batch of samples.

All samples were analyzed in positive (negative) polarity, using data-dependent acquisition mode. In this regard, full scan MS spectra (*m/z* 100–1700, scan rate 10 Hz, ID enabled) were followed by two data-dependent MS/MS spectra (*m/z* 100–1700, scan rate 10 Hz, ID disabled) for the two most intense ions per scan. The ions were selected when they reached an intensity threshold of 1500, isolated at the tuning file Q1 resolution, fragmented using collision induced dissociation (CID) with fixed collision energy (CE 20 eV), and excluded for 1 s before being re-selected for fragmentation. The parameters used for the ESI source were: interface voltage 4 kV ( -3 kV for negative polarity), interface temperature 300 °C, nebulizing gas flow 3 L/min, and drying gas flow 10 L/min. The parameters used for the CDS probe were: interface voltage 4.5 kV (positive polarity) or -3.5 kV (negative polarity), and nebulizing gas flow 1 L/min.

### SUPPLEMENTAL TABLES & FIGURES

#### Table S1. BioProjects which have more than one *P. syringae* strains harbouring the DHP cluster.

| BioProject | With DHP cluster | Total Assemblies | BioProject Description | Published in |
|---|---|---|---|---|
| PRJNA587608 | 39 | 171 | Epiphytic populations of Ps from cherry orchards. | 10.1111/nph.18573 |
| PRJNA750090 | 27 | 38 | Pseudomonas isolated from cherry trees from commercial orchards of Chile. | 10.1094/MPMI-04-22-0092-A 10.1094/PDIS-07-22-1638-PDN |
| PRJNA292453 | 17 | 268 | Pseudomonas Genome sequencing and assembly, comparative genome analysis of multiple strains of bacteria. | 10.1186/s13059-018-1606-y 10.1093/femsle/fnaa223 |
| PRJNA680595 | 17 | 58 | Collection of Pseudomonas strains that were isolated from infected plants in Turkey. | 10.1099/mgen.0.000585 |
| PRJNA345357 | 7 | 34 | Comparative genomics of P. syringae isolated from Prunus sp. and other hosts. | 10.1111/ppa.12834 |
| PRJNA728054 | 6 | 20 | P. syringae phylogroup 2 and P. viridiflava phylogroup 7 strains were isolated from bacterial stem blight-diseased alfalfa (Medicago sativa) plants from the Western and Midwestern United States. | Not yet. |
| PRJNA387592 | 6 | 18 | P. syringae isolated from A. thaliana from several population in the Midwestern USA. | 10.1371/journal.pone.0184195 |
| PRJNA481030 | 6 | 46 | Draft Genome Sequences of P. syringae Bean Isolates | Not yet. |
| PRJNA287460 | 4 | 26 | Genome sequencing and assembly of 26 lineages from the Pseudomonas syringae species complex. | 10.1111/mpp.12423 |
| PRJEB23287 | 2 | 9 | Prunus associated members of the P. syringae species complex | Not yet. |
| PRJNA396563 | 2 | 2 | P. syringae pv. syringae isolated from diseased cherry trees in South Africa | Not yet. |
| PRJNA603661 | 2 | 5 | Meat spoilage bacteria (Pseudomonas) | Not yet. |
| PRJNA274345 | 2 | 20 | Comparison of Pseudomonas species and description of novel species | 10.1099/ijsem.0.000852 10.1099/ijs.0.000230 |

#### Table S2. Plant metabolite elicitors used in Eco-HiTES screen.

| Elicitor | Vehicle | Final Concentration |
|---|---|---|
| Luteolin | DMSO | 20 µM |
| Homobrassinolide (28-Highbrassinolide) | DMSO | 20 µM |
| Dihydrojasmonic Acid | DMSO | 20 µM |
| *cis*-Zeatin | DMSO | 20 µM |
| Menthone | DMSO | 20 µM |
| Gibberellin A1 | DMSO | 20 µM |
| meta-Topolin | DMSO | 20 µM |
| Gibberellin A4 | DMSO | 20 µM |
| Nebularine | DMSO | 20 µM |
| *trans*-Zeatin Riboside | DMSO | 20 µM |
| Karrikinolide | DMSO | 20 µM |
| Gibberellic Acid | DMSO | 20 µM |
| Indole-3-butyric Acid | DMSO | 20 µM |
| Brassinolide | DMSO | 20 µM |
| (±)-Jasmonic Acid-Isoleucine | DMSO | 20 µM |
| N6-(Δ2-Isopentenyl)adenosine | DMSO | 20 µM |
| Epibrassinolide (24-Epibrassinolide) | DMSO | 20 µM |
| Indole-3-acetic Acid | DMSO | 20 µM |
| (+)-Abscisic Acid | DMSO | 20 µM |
| *trans*-Zeatin | DMSO | 20 µM |

| | | |
|---|---|---|
| (±)-Jasmonic Acid | DMSO | 20 µM |
| Jasmonic Acid methyl ester | DMSO | 20 µM |
| Indole-3-carboxylic Acid | DMSO | 20 µM |
| Glucosylceramide (from Soy, 2-hydroxy hexadecanoyl) | DMSO | 20 µM |
| β-Ocimene | DMSO | 20 µM |
| Acetosyringone | DMSO | 20 µM |
| Camalexin | DMSO | 20 µM |
| Brassinin | DMSO | 20 µM |
| Linamarin | DMSO | 20 µM |
| Concanavalin A (*Canavalia ensiformis*) | PBS (Sterile) | 20 µg/mL |
| A1-Phytoprostane-I | DMSO | 20 µM |
| 12-oxo Phytodienoic Acid | DMSO | 20 µM |
| Pterostilbene | DMSO | 20 µM |
| 13(S)-HpOTrE | DMSO | 20 µM |
| Palustric Acid | DMSO | 20 µM |
| Scopoletin | DMSO | 20 µM |
| Fraxetin | DMSO | 20 µM |
| Esculetin | DMSO | 20 µM |
| Apigenin | DMSO | 20 µM |
| Soyasaponin Bb | DMF | 20 µM |
| Timosaponin AIII | DMSO | 20 µM |
| Hederacoside C | DMF | 20 µM |
| Ginsenoside Re | DMSO | 20 µM |
| Platycodin D | DMSO | 20 µM |
| Glycyrrhizic Acid (ammonium salt) | DMSO | 20 µM |
| Isoastragaloside I | DMSO | 20 µM |
| Tenuifolin | DMSO | 20 µM |
| Notoginsenoside Ft1 | DMSO | 20 µM |
| Salicylic Acid | DMSO | 20 µM |
| Trigonelline (chloride) | water/ACN | 20 µM |
| Xanthurenic Acid | water/ACN | 20 µM |
| Syringic Acid | DMSO | 20 µM |
| Gangalin (3,5,7-trihydroxyflavone ) | DMSO | 20 µM |
| Hesperidin | DMSO | 20 µM |
| Vindoline | DMF | 20 µM |
| Indole 3 carbinol | DMSO | 20 µM |
| Cinnamic acid | water/ACN | 20 µM |
| *p*-Coumaric acid | water/ACN | 20 µM |
| Sinapic acid | water/ACN | 20 µM |
| 3-Hydroxycinnamic acid | water/ACN | 20 µM |
| Ferulic acid | water/ACN | 20 µM |

| | | |
|---|---|---|
| *p*-Hydroxybenzoic acid | water/ACN | 20 µM |
| Gallic acid | water/ACN | 20 µM |
| Vanillic acid | water/ACN | 20 µM |
| 4-Hydroxybenzaldehyde | DMSO | 20 µM |
| 3,4-Dihydroxybenzaldehyde | DMSO | 20 µM |
| Caffeic acid | water/ACN | 20 µM |
| Chlorogenic acid | DMSO | 20 µM |
| Coniferyl alcohol | DMSO | 20 µM |
| 4-Hydroxy-3-methoxycinnamaldehyde | DMSO | 20 µM |
| Syringaldehyde | DMSO | 20 µM |
| Myricetin | DMSO | 20 µM |
| Catechin | DMSO | 20 µM |
| Quercetin | DMSO | 20 µM |
| Morin | DMSO | 20 µM |
| Phytosphingosine | DMSO | 20 µM |
| L-Phenylalanine | water/ACN | 200 µM |
| L-Tyrosine | water/ACN | 200 µM |
| L-Tryptophan | water/ACN | 200 µM |
| L-Isoleucine | water/ACN | 200 µM |
| L-Leucine | water/ACN | 200 µM |
| L-Valine | water/ACN | 200 µM |
| Citrate | water/ACN | 200 µM |
| Fumarate | water/ACN | 200 µM |
| Malate | water/ACN | 200 µM |
| Succinate | water/ACN | 200 µM |
| Gaba (γ-aminobutyric acid) | water/ACN | 200 µM |
| L-Glutamic acid | water/ACN | 200 µM |
| L-Histidine | water/ACN | 200 µM |
| L-Ornithine | water/ACN | 200 µM |
| Thymidine | water/ACN | 200 µM |
| Uridine | water/ACN | 200 µM |
| L-Asparagine | water/ACN | 200 µM |
| L-Serine | water/ACN | 200 µM |
| L-Glutamine | water/ACN | 200 µM |
| L-Methionine | water/ACN | 200 µM |
| L-Alanine | water/ACN | 200 µM |
| Nicotinic Acid | water/ACN | 200 µM |
| Putrescine | water/ACN | 200 µM |
| Cytidine | water/ACN | 200 µM |
| Uracil | water/ACN | 200 µM |
| Orotic Acid | water/ACN | 200 µM |
| Shikimate Acid | water/ACN | 200 µM |

**Table S3. Oligonucleotides used in the study.**

| No. | Primer Name | Sequence(5'-3') |
|---|---|---|
| 1 | paaK-ko-u1-L | tcctccggtgatcgttcgcg |
| 2 | paaK-ko-u1-R-EcoRI | GTTGTAAAACGACGGCCAGTGAATTCactgcccatggtcttcgccgtc |
| 3 | paaK-ko-d2-R | gtcagctggaaggccgccc |
| 4 | paaK-ko-d2-L-HindIII | AGCTATGACCATGATTACGCCAAGCTTgcgctggtggagttcgcgaag |
| 5 | apra-paaK-ko-L-lu1L | gcacgcgaacgatcaccggaggaTCTAGAGGTGATGGATAACTTCG |
| 6 | apra-paaK-ko-R-ld2R | ccacccgggcggccttccagctgacTCTAGAGATGCGCGATAACTTCG |
| 7 | cDHP_A_F_pSET152_BamHI | tgggctgcaggtcgactctagaggatcctcgggcacggaacaagactgc |
| 8 | cDHP_A_R | tcctccggtgatcgttcgcgtg |
| 9 | cDHP_B_F | caccaccgcacacccgcac |
| 10 | cDHP_B_R | agcatggcggggctcttcct |
| 11 | cDHP_C_F | cgaggaaggaagagccccgc |
| 12 | cDHP_C_R_pSET152_EcoRI | ggaaacagctatgacatgattacgaattcaccctccggcgtcgtcttgt |

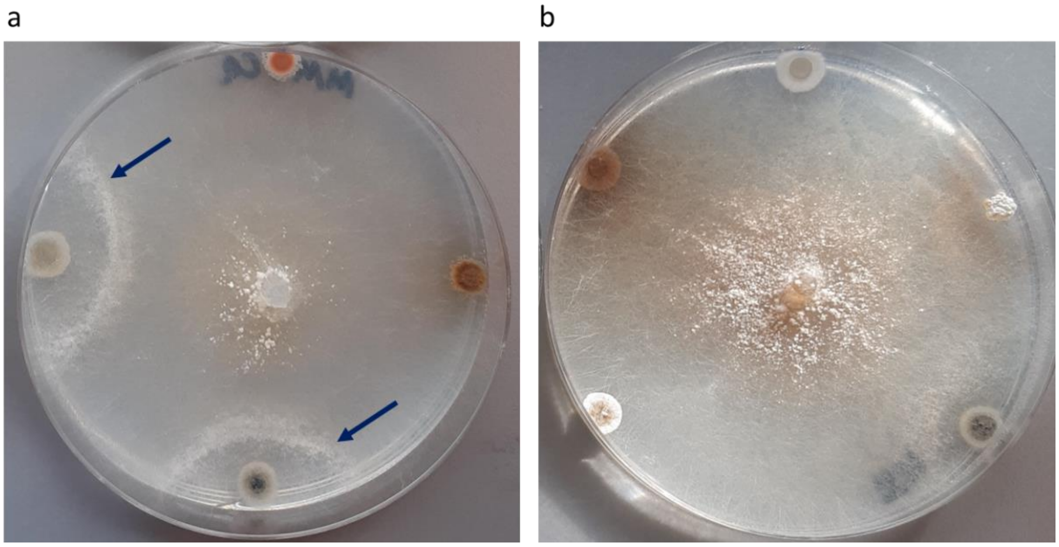


**Figure S1. Antifungal production by endophytic streptomycetes from the ATMOS collection.** Screening of endophytic streptomycetes isolated from *A. thaliana* for their ability to produce bioactive compounds inhibiting growth of *R. solani*. Bioactivity was assessed by growing isolates of the collection on the edge of a Petri dish on minimal medium. After five days, *R. solani* was inoculated in the middle of the plates. Inhibition of fungal growth was measured after a further three days of incubation. **a**) Five out of 35 strains inhibited hyphal growth of *R. solani*; arrows indicate strains that serve as examples; **b**) the remaining 30 did not show any bioactivity, with five representative inactive strains shown as an example.

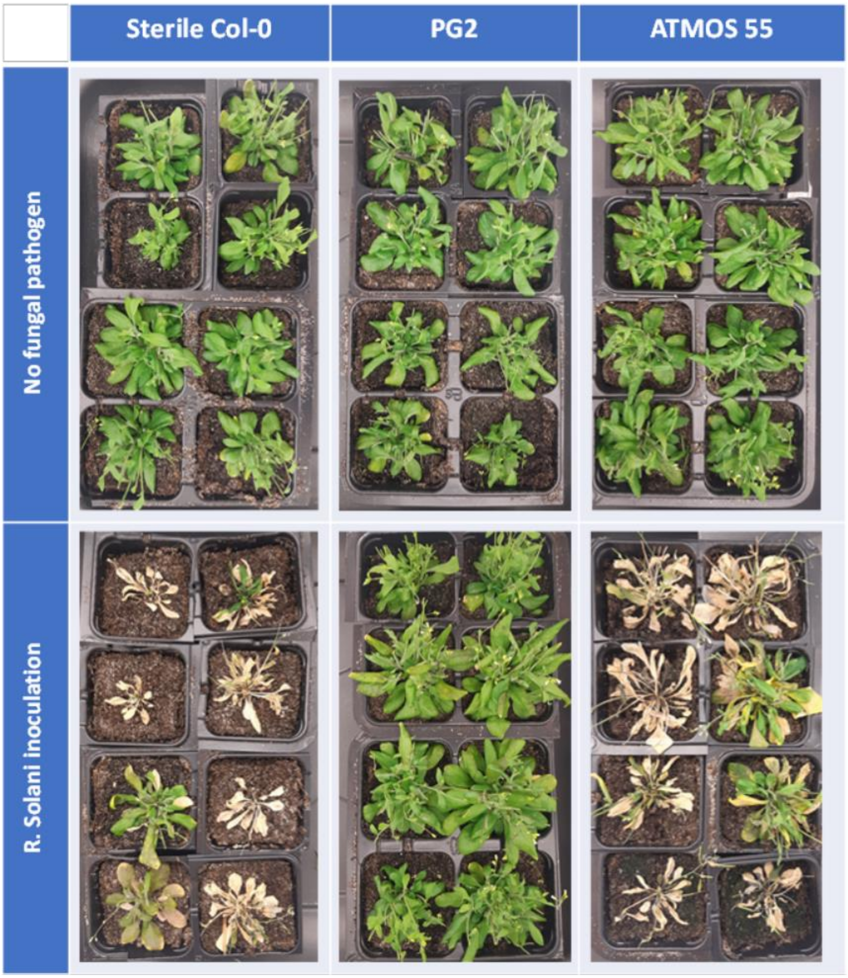


**Figure S2. In-soil evaluation of antifungal activity.** Top row: *A. thaliana* sterile or colonized by *Streptomyces*. Bottom row: the same, in the presence of *R. solani*. Only plants inoculated with *Streptomyces* sp. PG2 grew well in the presence of the fungal pathogen. Strains other than PG2 showing mild bioactivity against *R. solani* in the initial *in vitro* screening did not prevent the fungal infection. Plants colonized by *Streptomyces* ATMOS 55 are shown as an example.

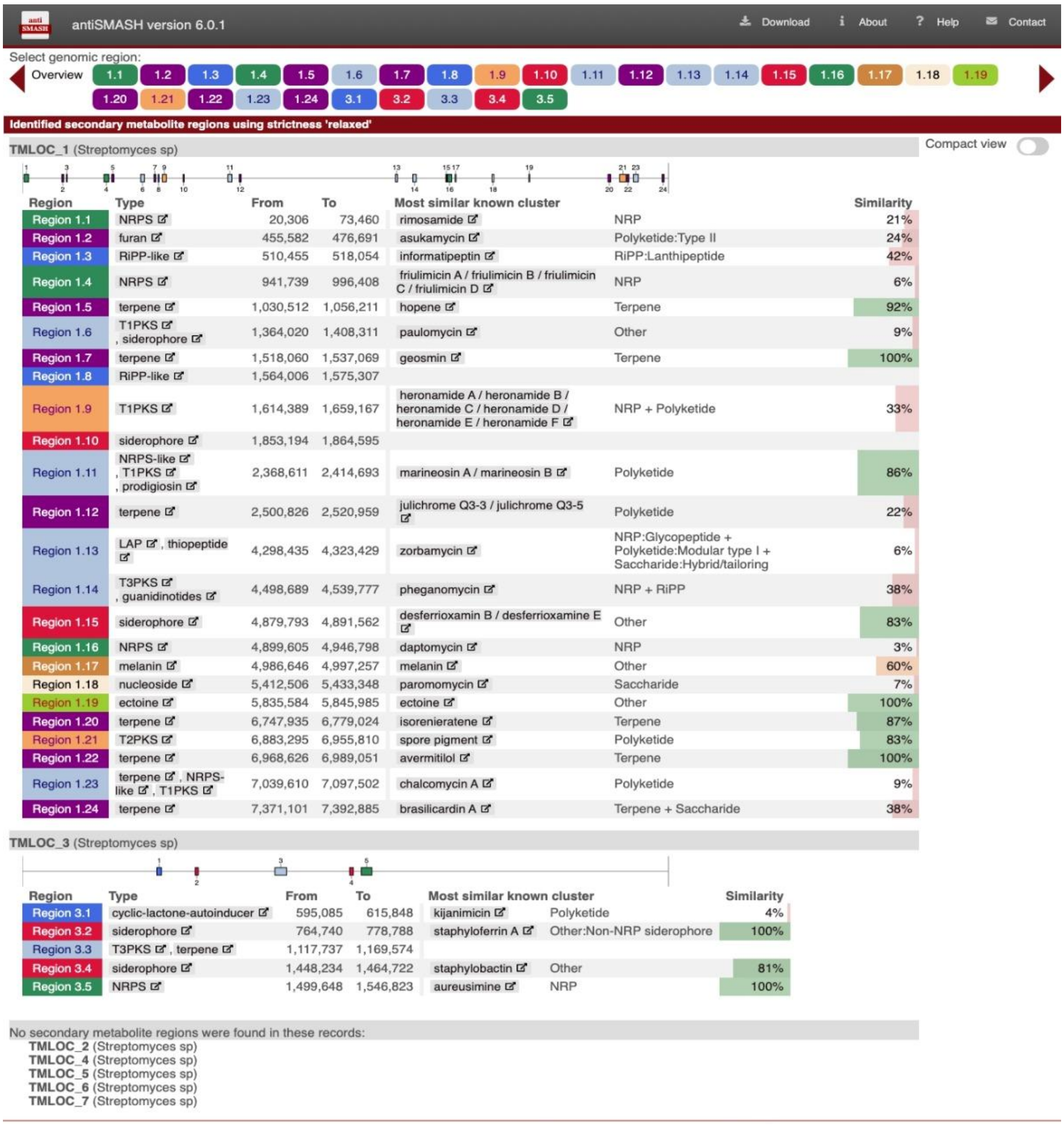
antiSMASH version 6.0.1 — Download — About — Help — Contact

Select genomic region:

Overview 1.1 1.2 1.3 1.4 1.5 1.6 1.7 1.8 1.9 1.10 1.11 1.12 1.13 1.14 1.15 1.16 1.17 1.18 1.19 1.20 1.21 1.22 1.23 1.24 3.1 3.2 3.3 3.4 3.5

Identified secondary metabolite regions using strictness 'relaxed'

TMLOC_1 (Streptomyces sp) — Compact view

| Region | Type | From | To | Most similar known cluster | | Similarity |
|---|---|---|---|---|---|---|
| Region 1.1 | NRPS | 20,306 | 73,460 | rimosamide | NRP | 21% |
| Region 1.2 | furan | 455,582 | 476,691 | asukamycin | Polyketide:Type II | 24% |
| Region 1.3 | RiPP-like | 510,455 | 518,054 | informatipeptin | RiPP:Lanthipeptide | 42% |
| Region 1.4 | NRPS | 941,739 | 996,408 | friulimicin A / friulimicin B / friulimicin C / friulimicin D | NRP | 6% |
| Region 1.5 | terpene | 1,030,512 | 1,056,211 | hopene | Terpene | 92% |
| Region 1.6 | T1PKS, siderophore | 1,364,020 | 1,408,311 | paulomycin | Other | 9% |
| Region 1.7 | terpene | 1,518,060 | 1,537,069 | geosmin | Terpene | 100% |
| Region 1.8 | RiPP-like | 1,564,006 | 1,575,307 | | | |
| Region 1.9 | T1PKS | 1,614,389 | 1,659,167 | heronamide A / heronamide B / heronamide C / heronamide D / heronamide E / heronamide F | NRP + Polyketide | 33% |
| Region 1.10 | siderophore | 1,853,194 | 1,864,595 | | | |
| Region 1.11 | NRPS-like, T1PKS, prodigiosin | 2,368,611 | 2,414,693 | marineosin A / marineosin B | Polyketide | 86% |
| Region 1.12 | terpene | 2,500,826 | 2,520,959 | julichrome Q3-3 / julichrome Q3-5 | Polyketide | 22% |
| Region 1.13 | LAP, thiopeptide | 4,298,435 | 4,323,429 | zorbamycin | NRP:Glycopeptide + Polyketide:Modular type I + Saccharide:Hybrid/tailoring | 6% |
| Region 1.14 | T3PKS, guanidinotides | 4,498,689 | 4,539,777 | pheganomycin | NRP + RiPP | 38% |
| Region 1.15 | siderophore | 4,879,793 | 4,891,562 | desferrioxamin B / desferrioxamine E | Other | 83% |
| Region 1.16 | NRPS | 4,899,605 | 4,946,798 | daptomycin | NRP | 3% |
| Region 1.17 | melanin | 4,986,646 | 4,997,257 | melanin | Other | 60% |
| Region 1.18 | nucleoside | 5,412,506 | 5,433,348 | paromomycin | Saccharide | 7% |
| Region 1.19 | ectoine | 5,835,584 | 5,845,985 | ectoine | Other | 100% |
| Region 1.20 | terpene | 6,747,935 | 6,779,024 | isorenieratene | Terpene | 87% |
| Region 1.21 | T2PKS | 6,883,295 | 6,955,810 | spore pigment | Polyketide | 83% |
| Region 1.22 | terpene | 6,968,626 | 6,989,051 | avermitilol | Terpene | 100% |
| Region 1.23 | terpene, NRPS-like, T1PKS | 7,039,610 | 7,097,502 | chalcomycin A | Polyketide | 9% |
| Region 1.24 | terpene | 7,371,101 | 7,392,885 | brasilicardin A | Terpene + Saccharide | 38% |

TMLOC_3 (Streptomyces sp)

| Region | Type | From | To | Most similar known cluster | | Similarity |
|---|---|---|---|---|---|---|
| Region 3.1 | cyclic-lactone-autoinducer | 595,085 | 615,848 | kijanimicin | Polyketide | 4% |
| Region 3.2 | siderophore | 764,740 | 778,788 | staphyloferrin A | Other:Non-NRP siderophore | 100% |
| Region 3.3 | T3PKS, terpene | 1,117,737 | 1,169,574 | | | |
| Region 3.4 | siderophore | 1,448,234 | 1,464,722 | staphylobactin | Other | 81% |
| Region 3.5 | NRPS | 1,499,648 | 1,546,823 | aureusimine | NRP | 100% |

No secondary metabolite regions were found in these records:

**TMLOC_2** (Streptomyces sp)
**TMLOC_4** (Streptomyces sp)
**TMLOC_5** (Streptomyces sp)
**TMLOC_6** (Streptomyces sp)
**TMLOC_7** (Streptomyces sp)

**Figure S3:** Prediction by antiSMASH of BGCs for secondary metabolites in the genome of *Streptomyces* sp. PG2. The DHP BGC was not predicted by the algorithm.

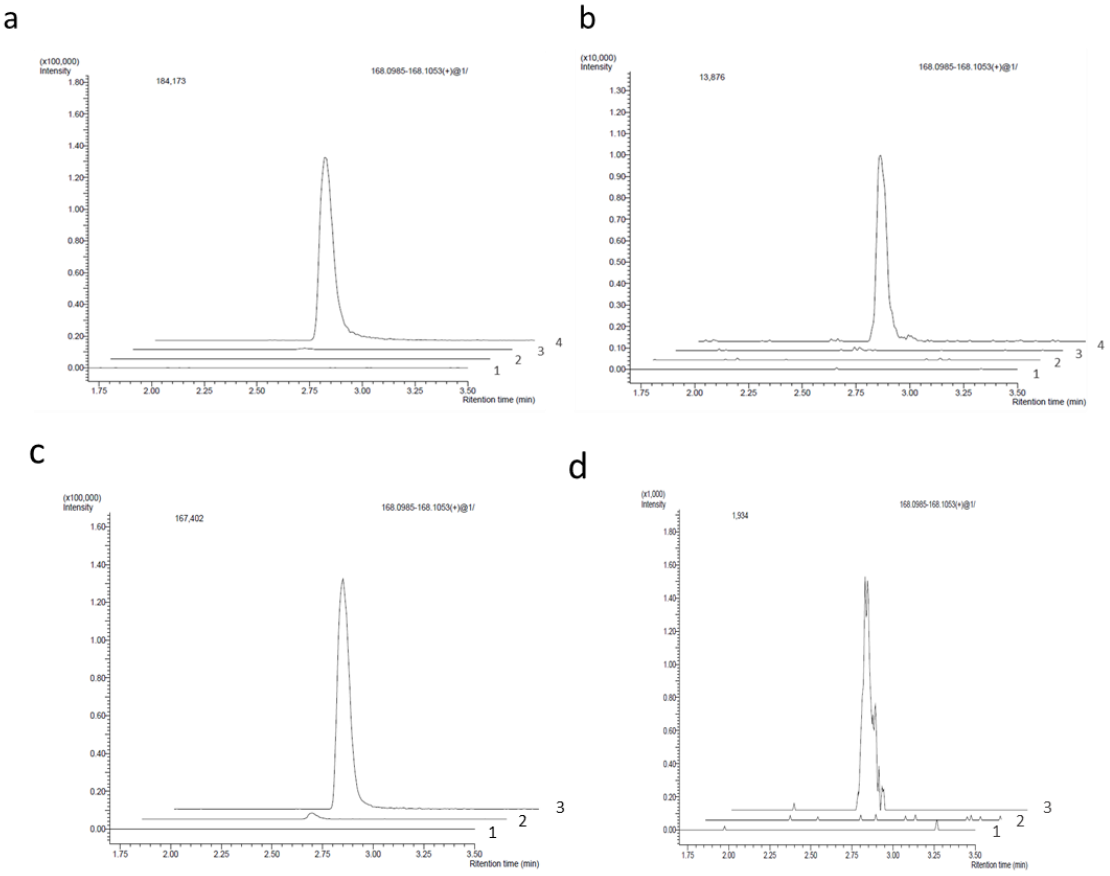


**Figure S4. Representative extracted ion chromatograms of DHP**. Chromatograms show DHP (168.1019 m/z) produced by bacteria and plants, as monocultures or in co-culture. **a) DHP production by the DHP null mutant and its complemented strain.** Mutant PG2_Δ*plu3041* in monoculture (curve 1) or in co-culture with *A. thaliana* seedlings (curve 2) failed to produce DHP. After genetic complementation with construct pE-DHP_Hyg containing the entire DHP BGC, DHP production was restored and could also be induced by co-cultivation with *A. thaliana*; this is shown by the low DHP production by the complemented mutant grown alone (curve 3), as compared to the strong productivity of the plant-microbe co-culture (curve 4). **b) Heterologous expression in *S. coelicolor* M145.** *S. coelicolor* M145 alone or in co-culture with *A. thaliana* seedlings (curves 1 and 2, respectively) failed to produce DHP. Introduction of a clone that allows heterologous expression of DHP in *S. coelicolor* led to a very limited production of DHP (curve 3); importantly, co-cultivation with *A. thaliana* strongly enhanced DHP production (curve 4), showing that all requirements for plant-dependent control are present in *S. coelicolor* and/or the DHP BGC. **c) Colonization of sugar beet seedlings also elicits DHP production.** Production of DHP was measured in extracts from sugar beet monoculture (curve 1), *Streptomyces* sp. PG2 monoculture (curve 2) or a co-culture of *Streptomyces* sp. PG2 with sugar beet seedlings (curve 3). Note that like with Arabidopsis, interaction with sugar beet also strongly elicited the production of DHP as compared to the bacterium grown alone. No DHP was produced by the plant monoculture. **d) Colonization-dependent elicitation of DHP production by *Pseudomonas syringae pv. syringae* FF5.** *Pseudomonas syringae pv. syringae* FF5 alone hardly produced DHP (curve 2), while strong production was observed in co-culture with *A. thaliana* (curve 3). No DHP was detected in plant monocultures (curve 1). This shows that also DHP production in *Pseudomonas* is induced by the plant.

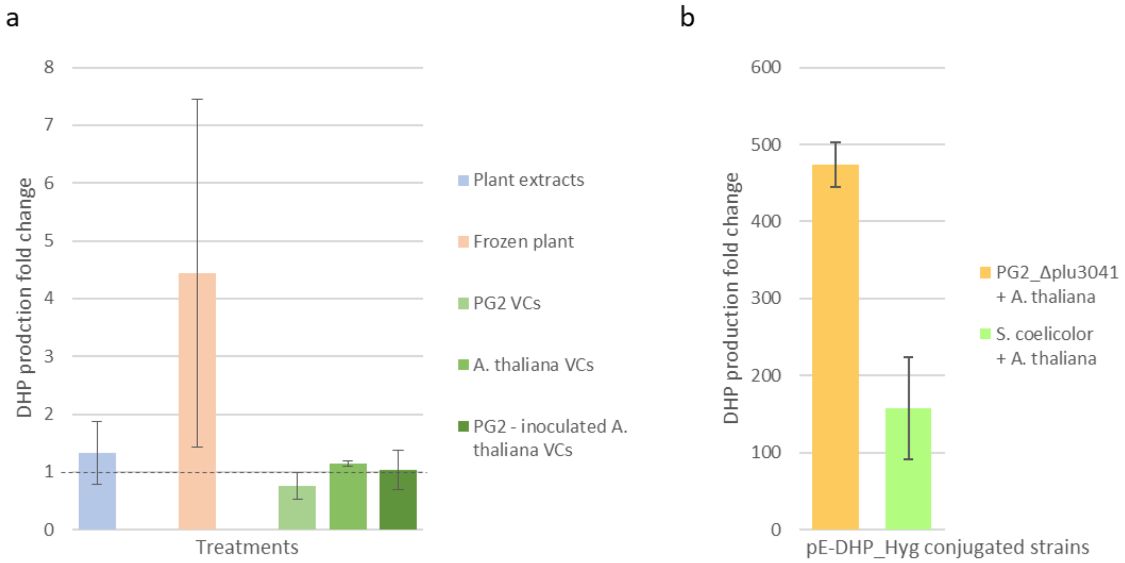


**Figure S5. DHP production fold change in different experiments.** Each bar represents one experiment performed in technical triplicate. **a) Investigation of possible elicitors of DHP in *Streptomyces* sp. PG2:** the streptomycete was cultivated in the presence of axenic plant spent plant medium (blue bar), dead plant material (red bar), or Volatile Compounds (VCs, green bars) from PG2 pure cultures, sterile A. thaliana or PG2-inoculated *A. thaliana*. None of these treatments resulted in significant increase in DHP production. **b) Heterologous expression of DHP BGC in PG2_Δ*plu3041* and *Streptomyces coelicolor* M145:** genetic complementation of the KO mutant **(orange bar)** restored the ability to produce DHP and increase its production upon plant signals; heterologous expression of DHP BGC in *S. coelicolor* confers to the strain the ability to produce DHP and to enhance its production when co-cultivated with *A. thaliana.*

a

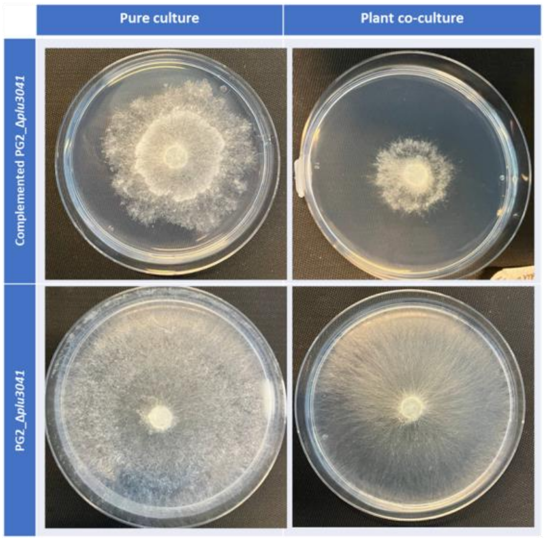


b

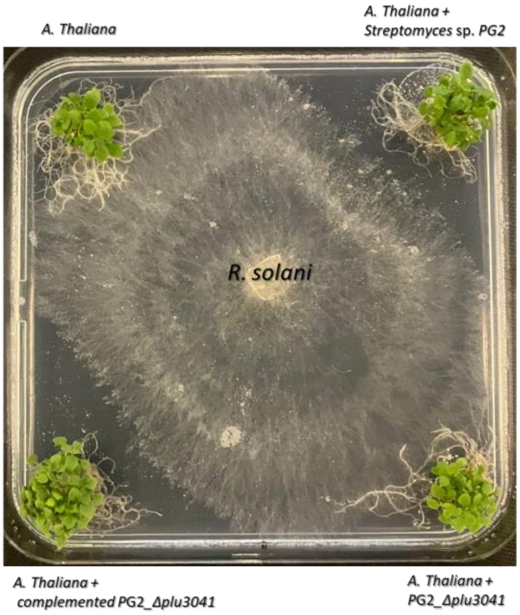


**Figure S6. Genetic complementation of mutant PG2_Δ*plu3041* restores the antifungal activity against *R. solani.* a)** In the top row, *R. solani* on plates containing spent media from genetically complemented PG2_Δ*plu3041*, grown either alone or in co-culture with *A. thaliana*, shows a plant-dependent increase in bioactivity against the fungal pathogen. In the bottom row, the same bioassay performed with the same strain without any complementation shows no antifungal activity. **b)** *R. solani*, growing from the centre of the plate, overgrows the axenic plant (top left) and the plant inoculated with PG2_Δ*plu3041* (bottom right). Genetic complementation of the DHP BGC restores the antifungal activity of the knock-out mutant, resulting in efficient plant protection (bottom left); plant protection by *Streptomyces* sp. PG2 wild-type strain is shown as a positive control (top right).

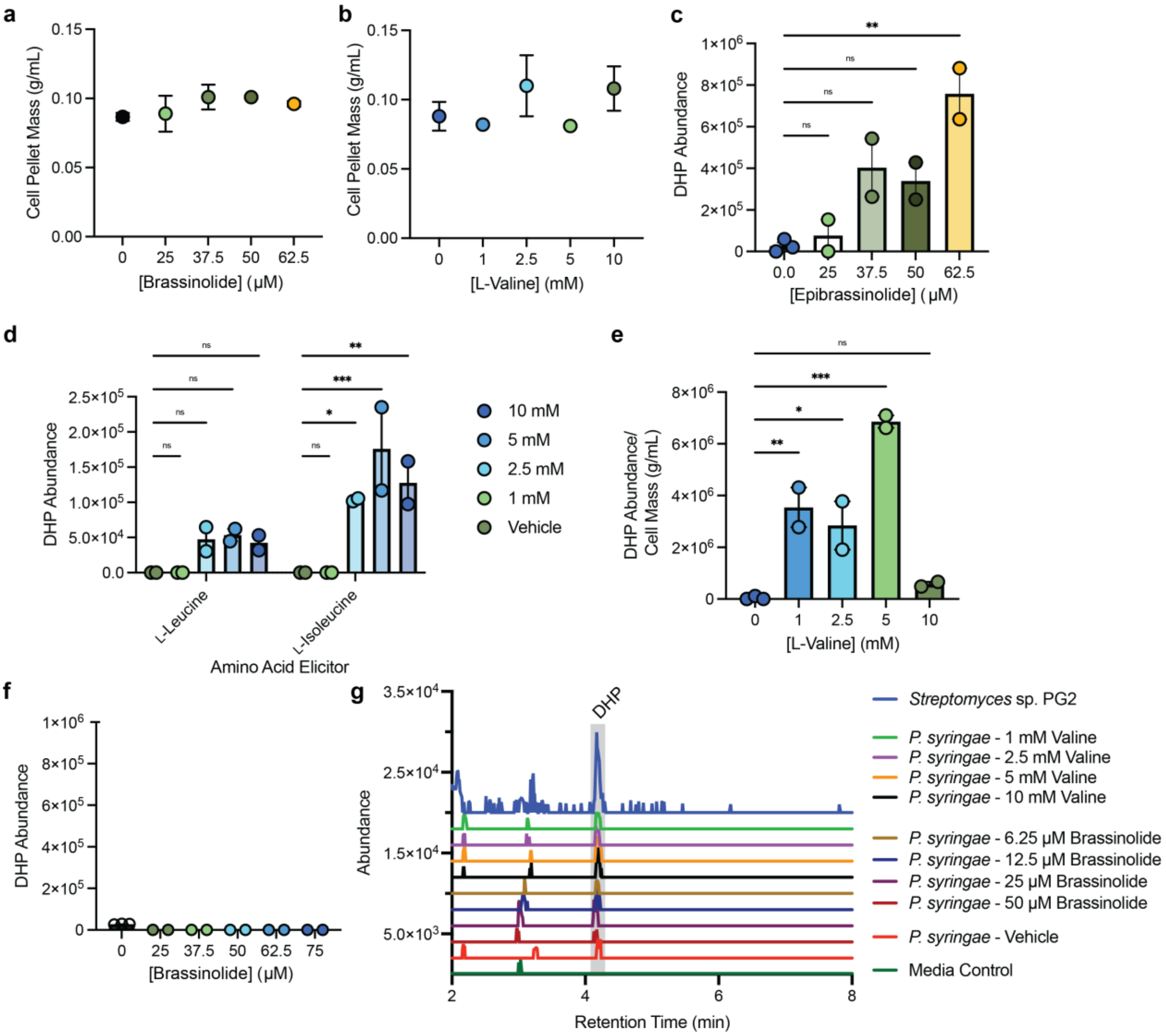


**Figure S7. Effects of brassinolides and BCAAs on DHP production and bacterial growth. a–b)** The effect of brassinolide **a)** and L-valine **b)** on growth of *Streptomyces* sp. PG2 determined by wet cell pellet mass. **c–d)** Elicitation of DHP in *Streptomyces* sp. PG2 by **c)** epibrassinolide and **d)** L-leucine and L-isoleucine. **e–f)** Elicitation, or lack thereof, of DHP in *S. coelicolor* harboring the DHP BGC with **e)** L-valine or **f)** brassinolide. **g)** EICs of 168.102 *m/z* from *P. syringae* at various concentrations of L-valine or brassinolide, indicating very low levels of DHP and no elicitation. An EIC of 168.102 *m/z* from *Streptomyces* sp. PG2 is shown to validate the retention time. For **a–f**, samples were grown as biological triplicates or duplicates, error bars represent the SE of the means. Significance for **c–e** was determined by Dunnett's multiple comparison test and for each, ***$P \leq 0.001$, **$P \leq 0.01$, *$P \leq 0.05$, & ns = not significant.

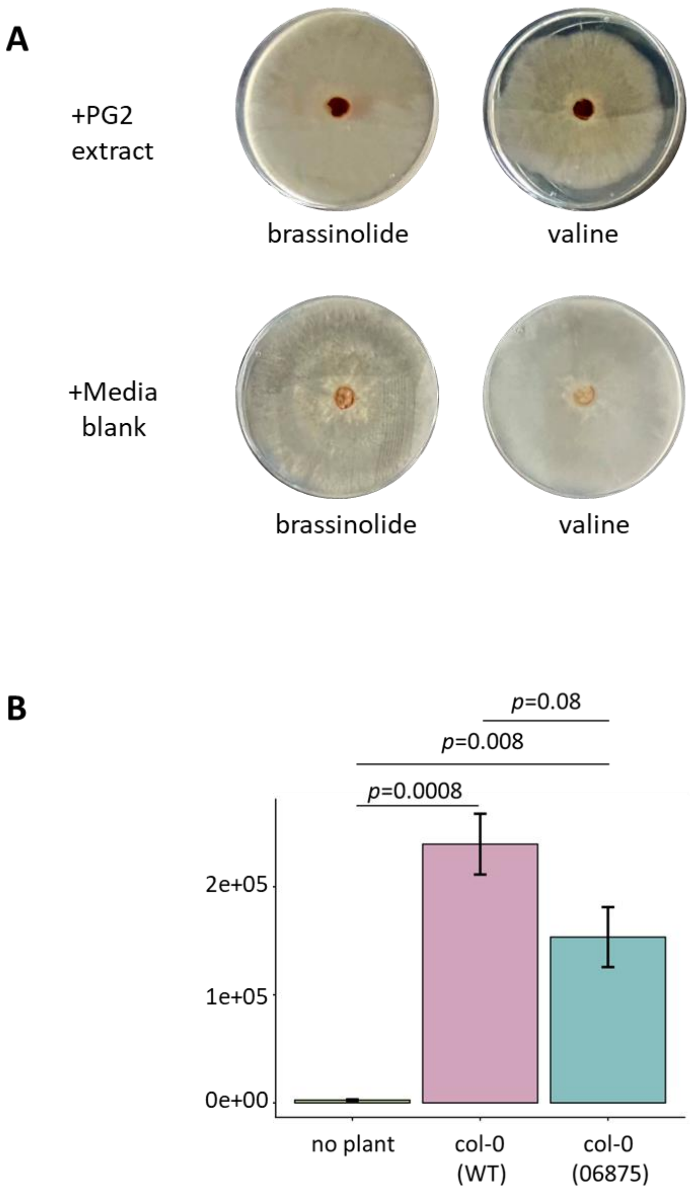

A
+PG2
extract
brassinolide
valine
none
+Media
blank
brassinolide
valine
B
p=0.08
p=0.008
p=0.0008
2e+05
1e+05
0e+00
no plant
col-0
(WT)
col-0
(06875)


**Figure S8.**
